# Coordinated Interaction: A model and test for globally signed epistasis in complex traits

**DOI:** 10.1101/2020.02.14.949883

**Authors:** Brooke Sheppard, Nadav Rappoport, Po-Ru Loh, Stephan J. Sanders, Andy Dahl, Noah Zaitlen

**Affiliations:** Department of Psychiatry and UCSF Weill Institute for Neurosciences, University of California, San Francisco; Bakar Computational Health Sciences Institute, University of California, San Francisco; Program in Medical and Population Genetics, Broad Institute of MIT and Harvard; Division of Genetics, Department of Medicine, Brigham and Women’s Hospital and Harvard Medical School; Departments of Neurology and Computational Medicine, University of California, Los Angeles

## Abstract

Interactions between genetic variants – epistasis – is pervasive in model systems and can profoundly impact evolutionary adaption, population disease dynamics, genetic mapping, and precision medicine efforts. In this work we develop a model for structured polygenic epistasis, called *Coordinated Interaction* (CI), and prove that several recent theories of genetic architecture fall under the formal umbrella of CI. Unlike standard polygenic epistasis models that assume interaction and main effects are independent, in the CI model, sets of SNPs broadly interact positively or negatively, on balance skewing the penetrance of main genetic effects. To test for the existence of CI we propose the *even-odd* (EO) test and prove it is calibrated in a range of realistic biological models. Applying the EO test in the UK Biobank, we find evidence of CI in 14 of 26 traits spanning disease, anthropometric, and blood categories. Finally, we extend the EO test to tissue-specific enrichment and identify several plausible tissue-trait pairs. Overall, CI is a new dimension of genetic architecture that can capture structured, systemic interactions in complex human traits.

## Introduction

Interaction between the phenotypic effects of genetic variants, or epistasis, is an essential component of biology with important consequences across multiple scientific domains. For example, epistasis significantly impacts evolutionary models, including response to selection or changing environment (Barton, 2017; Corbett-Detig et al., 2013). Epistasis also fundamentally shapes genetic architecture, as the direction of an allele’s effect can change based on genetic background (Park et al., 2018; Rau et al., 2019; Sittig et al., 2016). Epistatic interactions are pervasive in model systems, including in model organisms (Bloom et al., 2015; Brem et al., 2005; Corbett-Detig et al., 2013; Forsberg et al., 2017; Huang et al., 2012; Mackay, 2014) and in recent mammalian gene-level interaction screens (Dixit et al., 2016; Horlbeck et al., 2018; Norman et al., 2019). Moreover, these interactions often represent core biological functions. For example, the molecular chaperone HSP90 modifies diverse disease-model-specific proteins (Chen and Wagner, 2012; Meares et al., 2004; Queitsch et al., 2002), and conceptually similar mechanisms protect the cell from aberrant translation at ribosomes (Hickey et al., 2019). Such structured interactions are a core focus of systems biology, and their genetic bases have long been hypothesized to influence complex human disease.

While the observations in model organisms suggest that epistasis is also fundamental in humans (Mackay and Moore, 2014; Mackay, 2014), it remains poorly understood. This is largely because powerful, interpretable modelling tools are nascent. Genome-wide searches for interacting SNP pairs are computationally and statistically onerous, despite some success (Marchini et al., 2005). Additionally, recent methods for epistasis increase power by aggregating interactions across the whole genome (Crawford et al., 2017; Jannink, 2007; Park et al., 2018; Rau et al., 2019), and their results further support the potential importance of epistasis in complex traits. Interestingly, all of these approaches assume that epistasis is unstructured in that interaction effect sizes and directions are independent of main effects.

This is in contrast with recent conceptual models of complex human traits that imply or are consistent with structured forms of epistasis. For example, Zuk et al. describe a limiting pathway model of human disease that directly implies negative interactions between SNPs contributing to different pathways (Zuk et al., 2012). Also, the HSP90 community has discussed the possibility of a polygenic version of chaperon function (Milton et al., 2003), which would induce structured interactions between HSP90 buffer SNPs and exonic missense SNPs. Another example is the omnigenic model, which suggests that traits are determined by a few trait-specific “core” genes that are modified by many non-specific “peripheral” genes (Boyle et al., 2017; Liu et al., 2019). When these effects are non-linear, the omnigenic model implies structured epistasis between the SNPs contributing to “core” and “peripheral” genes.

Motivated by these concerns, we propose a model for structured epistasis, which we call *Coordinated Interaction* (CI). Conceptually, CI measures concerted interactions between pathways that are themselves additively heritable traits. We quantify CI with the parameter *γ*, where *γ* < 0 indicates negative epistasis between genetic pathways, on average, dampening the marginal effects of trait-increasing variants; conversely, *γ* > 0 indicates positive epistasis, where trait-increasing SNP effects are actually magnified in the full organismal context relative to their GWAS estimates.

For simplicity in exposition, we usually discuss examples with positive-valued traits, where positive/negative epistasis coincide with the more interpretable notions of synergy/antagonism; in general, however, the skew in genetic risks driven by *γ* ≠ 0 may either dampen or exacerbate marginal genetic effects (Phillips, 2008). In Figure 1, we illustrate *γ* in the case of two latent interacting pathways under the positive phenotype assumption. The key is that *γ* ≠ 0 if there are heritable, positively (synergistically) or negatively (antagonistically) interacting pathways (Figure 1b), while *γ* = 0 if the interactions are purely random or absent (Figure 1a).

**Figure 1.**
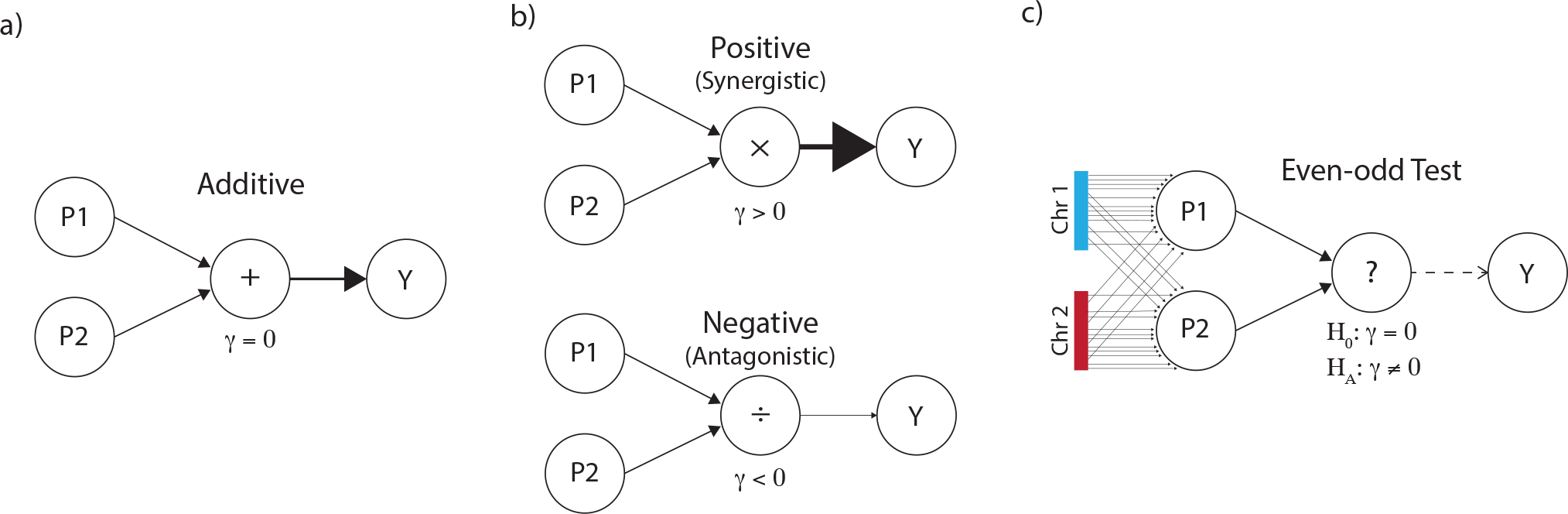
Coordinated Interaction and the Even-Odd test with two chromosomes. (a) In the additive model SNP effects from the two pathways are summed to produce the positive phenotype (*γ* = 0). (b) Same as (a), except the pathways interact either positively (synergistically, ×, *γ* > 0) or negatively (antagonistically, ÷, *γ* < 0). (c) The even-odd test considers interaction from traits derived from the even and odd chromosomes in place of the unknown pathways truly driving the interaction.

Testing for the existence of CI would be trivial if we knew which SNPs contributed to which pathways: we could simply build genetic predictors of each pathway and test their interaction. However, these pathways are generally unknown, making CI estimation seem impossible. Surprisingly, we show that testing for CI can be accomplished by randomly assigning independent sets of SNPs to arbitrary pathways. Concretely, we build polygenic risk scores (PRS) specifically for the even and odd chromosomes and then test their interaction in a linear regression on phenotype (Figure 1c). We call this the *even-odd* (EO) test and we prove analytically that EO reliably estimates CI under polygenicity. We also prove that under a polygenic model, any chromosome split, not just even-odd, gives identical results in expectation. Importantly, however, partitions aligned closely with the true latent pathways will have greater power. We leverage this fact to test for CI enrichment in specific genomic annotations. Specifically, we focus on tissue-specific annotations to ask whether complex traits are enriched for interactions between tissue-specific pathways.

In this paper, we first introduce the concept of Coordinated Interaction (CI) formally and define the even-odd (EO) test. We then examine simulated data sets with assortative mating and population structure to show that EO is robust to confounding. Then we perform the EO test in the UK Biobank (UKBB) across 26 traits spanning multiple domains and find 14 with Bonferroni-significant CI. We validate the approach, which uses internally cross-validated PRS, by using PRS constructed from external data sources for 8 of the 26 traits. Finally, we estimate tissue-specific CI across thirteen tissues in the UKBB and find several biologically plausible tissue-trait pairs, as well as enrichment for interacting tissue pairs. We conclude with a discussion of limitations to our approach, implications for association testing and genetic architecture, and possible future extensions.

## Results

### Coordinated Polygenic interaction

Throughout the paper, we assume a polygenic pairwise interaction model:

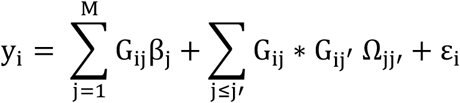

y_i_ is the phenotype for individual *i*, and G_ij_ is the genotype for individual *i* at marker *j*. *ε* ∼ *i*.*i*.*d*. *N*(0, *σ*^2^) is the residual error. The vector β contains the marginal polygenic effects. Ω*_jj_*, is the pairwise interaction effect of SNPs j not equal to j′, so Ω is the matrix of all pairwise SNP interaction effects in the genome.

The standard additive model assumes no epistasis, i.e. Ω = 0. In this model, SNP *j* always has the same effect β_j_, on the phenotype, regardless of genetic background or environment. In the polygenic setting, where there are many more SNPs than individuals (M > N), total heritability can still be reliably estimated by the random-effect model GREML, which models β_j_, as i.i.d. Gaussian (Yang et al., 2010)

Epistatic models go further by allowing nonzero Ω. To date, epistatic tests have focused either on candidate SNPs or genome-wide screens for SNP pairs, which reduce M<N and facilitate simple fixed-effect models (Marchini et al., 2005). More recently, random-effect models akin to GREML have become popular for estimating the total size of Ω (i.e. the heritability from pairwise epistasis) (Cockerham, 1954; Henderson, 1985; Jiang and Reif, 2015; Young and Durbin, 2014). Another recent approach tests for interaction between a single SNP and a genome-wide kinship matrix, a useful compromise that provides SNP-level resolution and also aggregates genome-wide signal (Crawford et al., 2017).

While these methods are useful for characterizing the existence and impact of epistasis, all are limited by the assumption that β and Ω are independent. We are interested in an orthogonal question: When are β and Ω deeply intertwined by latent interacting pathways? Conceptually, β and Ω encode all relevant information, so the goal is to decode the presence of interacting pathways from these parameters. Concretely, we prove that these pathways exist if and only if the Coordinated Interaction γ is nonzero, where γ is defined as:

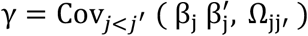

Intuitively, γ < 0 negatively skews the total polygenic effect relative to additivity. In the case that phenotypes are positive, this is equivalent to dampening the aggregate additive heritability, or antagonism between main effects, and it skews the population of phenotypes toward 0. Conversely, γ > 0 positively skews the population, increasing the probability of extremely high phenotypes. Note that γ = 0 does not imply that interactions are absent; rather that interactions do not necessarily live in the space of pathways that have main effects, which is exactly the model assumed by the above uncoordinated epistatic models.

For example, imagine there are two genetically independent pathways that are each sufficient for T2D, one based on BMI and one based on pancreas function. Then γ < 0 is expected, because high-BMI cases are not likely to also have high pancreas risk, which is rare. On the other hand, if a disease requires impairment across multiple distinct systems—e.g. if asthma requires both immune components and environmental exposures—then γ > 0 is expected: the impact of a smoking SNP on asthma is much larger if the immune system is compromised.

We provide a more rigorous exploration of coordination in the Supplementary Material. In particular, we prove that several biologically plausible interaction models induce CI, including the polygenic generalization of molecular buffers (Chen and Wagner, 2012), limiting disease pathways (Zuk et al., 2012), *trans* genetic regulation (Liu et al., 2019), and gene-environment interaction with heritable environment (Dahl et al., 2020).

### The Even-Odd estimator for Coordinated Interaction

We have defined our target, the Coordinated Interaction γ, as a function of the true genetic effects β and Ω. However, these parameters are not known; even worse, they are high-dimensional and cannot be accurately estimated. Testing for nonzero γ seems hopeless.

Our key idea in this paper is that randomly defined pathways can act as proxy for the true pathways. These random pathways will interact if and only if there are latent pathways that truly interact. Fundamentally, this is because the random approach appropriately sorts many pairs of SNPs into the right partition. Under true interacting pathways, these SNP pairs will drive interactions between the arbitrary PRS. Although the arbitrary pathways will miss some signal—from interacting pairs that are incorrectly placed in the same pathways—this does not cause false positives and, moreover, can be corrected *post hoc* under an infinitesimal model.

We propose estimating γ by regressing on the interaction between PRS built specifically from even and odd chromosomes (PRS_*e*_ and PRS_*o*_):

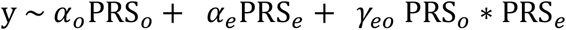

The ordinary least squares estimate 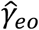 is the Even-Odd estimator of the coordination *γ*. We prove that *γ*_*eo*_ = 0 if and only if *γ* = 0, so we can simply use a regression test for 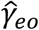 to test for the existence of CI. We describe the test formally in the Supplementary Material, including details of PRS construction, covariate adjustment, and modelling assumptions.

Under an infinitesimal model, all random, independent SNP sets give identical CI results. With finite genomes, instead, we can evaluate several partitions to maximize power. It is important that SNPs are independent across pathways to avoid false positives from nonlinear per-variant epistasis, hence we always evaluate chromosome-level genome partitions. This is analogous to our choice to exclude the Ω_jj_ terms from CI: in both cases, CI implies nonlinear effects of each individual SNP and the aggregate PRS, but the converse is not true. We use Bonferroni correction to adjust for multiple testing across multiple chromosome splits, but less conservative tests may add substantial power in the future as the EO test is highly correlated across different chromosome splits.

### Even-Odd distinguishes coordination from population structure, assortative mating, and uncoordinated interactions in simulation

To examine the properties of the EO test, we simulate data and examine the PRS and *γ*_*eo*_ under three biologically plausible genetic architectures: additivity, uncoordinated (i.e. random) interaction, and Coordinated Interaction. For each architecture, we simulate either unconfounded genotypes and phenotypes, or add confounding population structure or assortative mating (see Methods). Together, these settings represent a realistic range of causal and non-causal components of the genotype-phenotype map (Table 1).

**Table 1.** Polygenic simulations under additivity, uncoordinated interaction, or coordinated interaction assuming random mating, population structure, or assortative mating. Mean, standard deviation, and positive rate are shown for estimates of *θ_eo_* and *γ_eo_* for 10,000 simulations, where all SNPs were randomly assigned to either PRS_*e*_ or PRS_*o*_. PR is the positive rate, or the proportion of significant test statistics at *α* = 0.05; sd is standard deviation.

| | Population Characteristics | | $\theta_{eo}$: $\text{PRS}_e \sim \text{PRS}_o$ | | | | $\gamma_{eo}$: $Y \sim \text{PRS}_e + \text{PRS}_o + \text{PRS}_e * \text{PRS}_o$ | | | |
| --- | --- | --- | --- | --- | --- | --- | --- | --- | --- | --- |
|  | Assortative Mating | Population Structure (FST = 0.1) | -PCs |  | +PCs |  | -PCs |  | +PCs |  |
| | | | $X^2$ (sd) | PR $\alpha=0.05$ | $X^2$ (sd) | PR $\alpha=0.05$ | $X^2$ (sd) | PR $\alpha=0.05$ | $X^2$ (sd) | PR $\alpha=0.05$ |
| Additive | 0 | 0 | 1.0 (1.4) | 0.0487 | -- | -- | 1.0 (1.4) | 0.0469 | -- | -- |
|  | 1 | 0 | 26.4 (10.2) | 0.9992 | -- | -- | 1.0 (1.4) | 0.0496 | -- | -- |
|  | 0 | 1 | 7.6 (16.7) | 0.3343 | 1.0 (1.4) | 0.0502 | 1.0 (1.4) | 0.0498 | 1.0 (1.4) | 0.0504 |
| Uncoordinated Interaction | 0 | 0 | 1.0 (1.4) | 0.0483 | -- | -- | 1.0 (1.4) | 0.0491 | -- | -- |
|  | 1 | 0 | 17.7 (8.3) | 0.9845 | -- | -- | 1.0 (1.4) | 0.0513 | -- | -- |
|  | 0 | 1 | 7.5 (17.7) | 0.3354 | 1.0 (1.4) | 0.0519 | 1.0 (1.4) | 0.0489 | 1.0 (1.4) | 0.0507 |
| Coordinated Interaction | 0 | 0 | 1.0 (1.4) | 0.0458 | -- | -- | 18.7 (17.5) | 0.7537 | -- | -- |
|  | 1 | 0 | 14.7 (10.1) | 0.839 | -- | -- | 55.4 (50.0) | 0.8523 | -- | -- |
|  | 0 | 1 | 7.8 (17.7) | 0.3345 | 1.0 (1.4) | 0.0474 | 13.7 (13.9) | 0.6884 | 14.9 (14.6) | 0.712 |

First, we regressed PRS_*e*_ on PRS_*o*_ to calculate their correlation *θ*_*eo*_, which is related to an existing estimate of assortative mating (Yengo et al., 2018). We found that the test for *θ*_*eo*_ ≠ 0 reliably indicated the presence of assortative mating or uncorrected population structure. Note that after adjusting for PCs the test for *θ*_*eo*_ ≠ 0 was roughly null in the presence of population structure.

Our main focus, though, is on the test for *γ*_*eo*_ ≠ 0. We find that the null is well-behaved both under pure additivity and under uncoordinated pairwise interaction, with false positive rates near .05 at a p<.05 threshold. The EO test has much higher signal under CI (power >70%), showing the test has power to detect true CI. Importantly, neither uncorrected population structure nor assortative mating induced false positives for the EO test. Nevertheless, in practice, we recommend adjusting for population structure when running the EO test.

The power of the EO test is partially reduced by the fact that SNPs contributing to each of the latent interacting pathways are randomly distributed across the even and odd chromosomes (Supplementary Material). When we instead constructed PRS based on the true latent pathways, power to detect CI increased substantially, but the test was similar in all other respects (Supplementary Table 1). This shows that the EO test power indicates the accuracy of the partition used to construct PRS, increasing as the partition homes in on the true pathways.

### Identifying traits with Coordinated Interaction in UK Biobank

We next tested for Coordinated Interaction with the EO test in the UK Biobank (UKBB) (Bycroft et al., 2018). We studied 21 quantitative and 5 binary traits (Supplementary Table 2) chosen to represent a range of trait classes including anthropometric, disease, and blood traits. We specifically analyzed the subjects classified as “White British” and filtered out related individuals to minimize population structure bias while retaining large sample sizes (max n = 393076; Supplementary Table 2). We calculated PRS for each chromosome for each sample using a standard clumping + thresholding approach (Euesden et al., 2015) (Methods). The total PRS is then the sum of the individual chromosomes’ PRS. Thresholds were chosen to maximize percent variance explained (PVE) by the total PRS (Methods). We used 10-fold cross validation to minimize bias from in-sample overfitting. We summarize these cross-validated PRS in Table 2 and show the variability of across chromosomes in Supplementary Table 2.

**Table 2.** Even-odd test results for 12 traits with significant Coordinated Interaction in the UK Biobank. Significant traits, and external PRS replication for significant internal traits, are shown; results for all traits are shown in Supplementary Table 2. We summarize evidence for CI per trait using the minimum p-value over 100 random bifurcations, and we provide the CI estimate (γ) for this top split. *τ* is the marginal SNP p-value threshold used to construct the PRS and is chosen to maximize cross-validated prediction accuracy. “Int” indicates that the PRS was calculated using the cross-validation method with UKBB data (Methods). “Ext” indicates that the PRS was calculated using GWAS summary statistics from external datasets (Methods). PRS %VE describes the variance explained by the chosen PRS across all samples. * indicates p<0.05/100; ** indicates p<.05/100/#tissues; and *** indicates p<.05/100/#tissues/#phenotypes

| Phenotype | PRS Type | $\tau$ | Signif. Level | Min p | $\gamma$ | PRS %VE |
| --- | --- | --- | --- | --- | --- | --- |
| Platelet Vol. | Int Cont | 0.01 | ** | 2.60E-18 | -1.60E-03 | 20.7 |
| Platelet # | Int Cont | 0.01 | ** | 1.30E-05 | -9.40E-04 | 12.6 |
| Height | Int Cont | 0.1 | * | 1.10E-04 | -3.00E-04 | 11.7 |
| Platelet Distn Width | Int Cont | 0.01 | * | 1.10E-04 | -5.50E-04 | 9.3 |
| Sphered Cell Vol | Int Cont | 0.01 | * | 1.80E-04 | -1.20E-03 | 9.2 |
| Monocyte # | Int Cont | 0.01 | ** | 1.40E-05 | -6.20E-04 | 7.4 |
| LDL | Int Cont | 0.0001 | ** | 7.30E-11 | -8.20E-03 | 5.9 |
| Edu Years | Int Cont | 1 | ** | 2.10E-06 | 4.10E-04 | 5.7 |
| BMI | Int Cont | 1 | ** | 1.80E-06 | -3.00E-04 | 5.6 |
| Lymphocyte # | Int Cont | 0.1 | * | 4.40E-04 | -3.40E-04 | 5.5 |
| Eosinophil # | Int Cont | 0.01 | * | 3.20E-04 | -4.20E-04 | 5.0 |
| Triglycerides | Int Cont | 0.01 | * | 2.50E-04 | -1.30E-03 | 4.8 |
| T2D | Int Bin | 0.001 | * | 3.30E-05 | -5.00E-03 | 0.7 |
| Asthma | Int Bin | 0.001 | ** | 2.50E-07 | 5.60E-03 | 0.6 |
| Edu Years | Ext Cont | 1 | ** | 7.40E-07 | 1.50E-06 | 7.7 |
| Height | Ext Cont | 1 |  | 2.90E-03 | 8.00E-07 | 7.3 |
| BMI | Ext Cont | 1 | * | 3.70E-04 | -2.60E-06 | 5.6 |
| Triglycerides | Ext Cont | 0.01 | * | 5.50E-05 | -1.10E-05 | 3.5 |
| LDL | Ext Cont | 0.01 | ** | 7.70E-08 | -2.40E-05 | 2.6 |
| T2D | Ext Bin | 1 |  | 4.90E-02 | -1.30E-05 | 0.7 |
| Asthma | Ext Bin | 0.0001 | * | 2.90E-04 | 8.10E-05 | 0.6 |

Unlike in perfectly infinitesimal models, in real data the EO test depends on the specific bifurcation of chromosomes used to estimate *γ*. To minimize bias from choosing a single split, e.g. even vs. odd, in practice we analyzed CI estimates from 100 different random partitions of the 22 autosomes into two groups. To jointly analyze all bifurcations, we used Bonferroni adjustment, which is highly conservative as these tests are tightly correlated—taking height, for example, the correlation between the interaction terms (PRS_*o*_ * PRS_*e*_) is roughly 50% across splits, as expected under minimal inter-chromosomal LD (IQR is [48%,56%]). Following common practice (Finucane et al., 2018, 2015; Justice et al., 2019; Loh et al., 2018; Schoech et al., 2019; Zhu et al., 2018), we report the significant results per phenotype. The figures also denote the Bonferroni threshold over all splits and phenotypes, which is additionally overly conservative because of the high correlation between phenotypes.

Despite this aggressively conservative multiple testing adjustment, we discovered 14 traits with significant Coordinated Interaction, including T2D, asthma, height, educational attainment (EA), and BMI (Figure 2). We also detected CI for 9 blood traits: mean platelet volume (MPV), platelet distribution width (PDW), sphered cell volume (SCV), low density lipoprotein (LDL), triglycerides (TG), and counts for platelets (PLAT), monocytes (MONO), lymphocytes (LYM), and eosinophils (EOS) (Figure 2). The CI estimates themselves revealed trait-specific patterns of positive/negative epistasis (Table 2, Supplementary Table 2, Supplementary Figure 1). For MPV, PDW, PLAT, height, SCV, LDL, TG, LYM, and BMI, every bifurcation yielded negative estimates of *γ*, suggesting the latent pathways generally interact negatively; also, 86%, 90%, and 99% of EOS, MONO, and T2D splits had *γ* < 0. Conversely, asthma and EA had positive estimates for 90% and 100% of bifurcations, respectively, suggesting ‘and’ logic predominates across systems—e.g. stylistically, asthma results only if multiple systems fail. Importantly, these signs are invariant under modest phenotype rescaling (Supplementary Material). Taken together, these results show that Coordinated Interactions contribute the genetic architecture of multiple complex traits.

**Figure 2.**
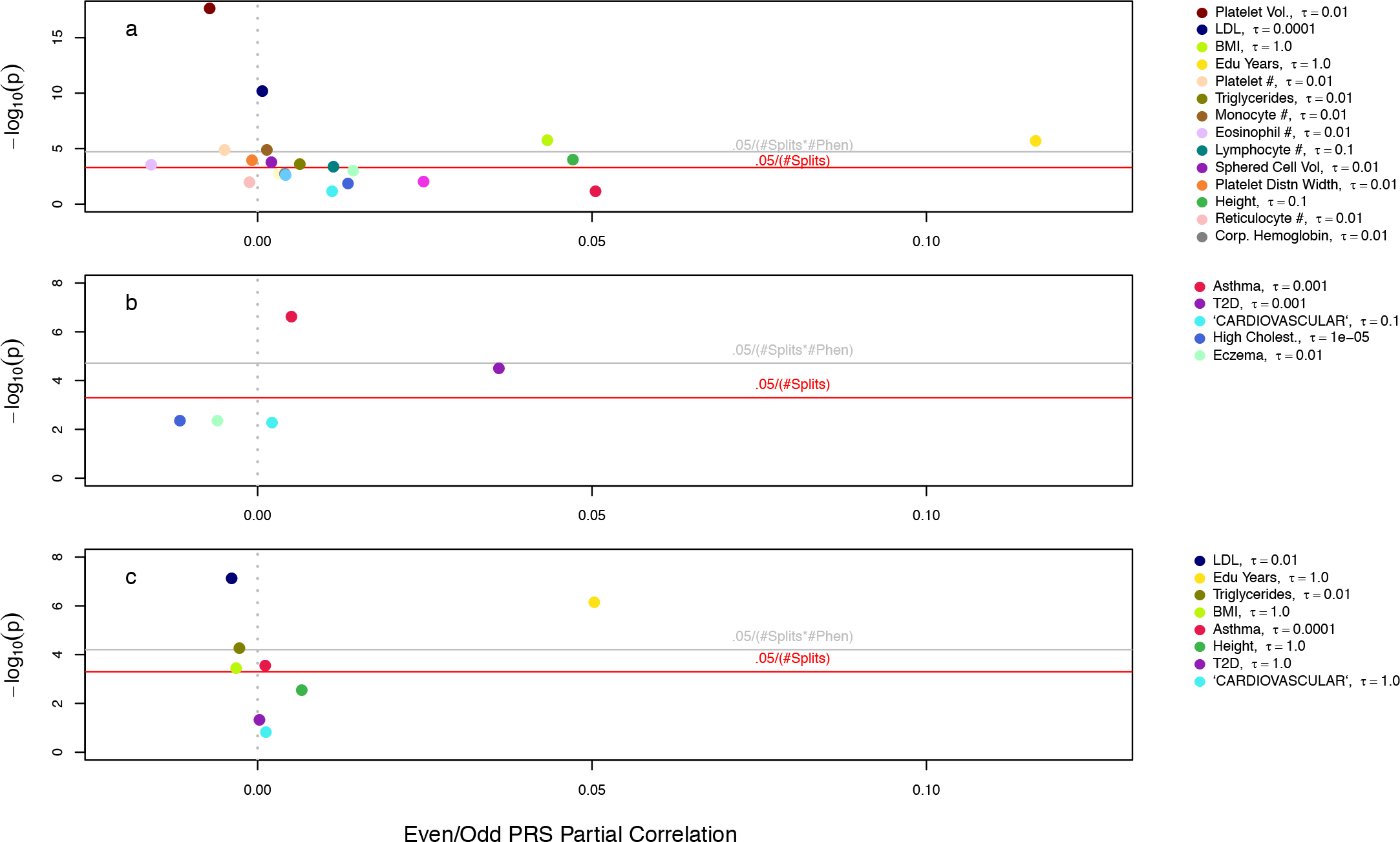
Coordinated Interaction in the UK Biobank. Minimum p-value split for EO test using cross-validated in-sample PRS for 21 quantitative traits (a) and 6 binary traits. (c) EO tests using external PRS for 8 of the quantitative and binary traits in (a) and (b) with external GWAS summary statistics available. In (a), the legend is only provided for the traits with highest mean –log10(p). *τ* is the p-value threshold used to construct the PRS and is chosen to maximize cross-validated prediction accuracy.

To assess the potential for impact by population structure and assortative mating, we calculate the correlation between the even and odd PRS (*θ*_*eo*_). Similar to (Yengo et al., 2018), we found that EA and height had substantial *θ*_*eo*_. Nonetheless, we observed no relationship between the *θ*_*eo*_ and *γ*_*eo*_ across splits within phenotypes. Furthermore, we saw no inflation of the EO test in simulation even when population structure was uncorrected and assortative mating was present. We conclude that there is little evidence for confounding of the EO interaction test results by uncorrected structure or assortative mating.

### Replicating Coordinated Interaction with external PRS in UK Biobank

Although we have cross-validated our PRS, we wish to additionally check that our results are not specific to a single population by overfitting dataset-specific confounders (Berg et al., 2019; Kerminen et al., 2019; Sohail et al., 2019). We investigated this by constructing PRS using external summary statistics to remove the potential impact of cohort specific artifacts. We selected eight traits that have large external GWAS summary statistics: asthma, T2D, cardiovascular disease, height, BMI, TG, EA, and LDL.

Our CI results replicated in sign and significance for asthma, BMI, TG, EA, and LDL (Table 2). However, significant CI did not replicate for height or T2D. The discrepancy for height may arise from well-known confounding issues in the GIANT summary statistics we used (Berg et al., 2019; Sohail et al., 2019). The T2D non-replication likely reflects loss of power, on the other hand, as 93% of splits replicated the negative *γ* < 0 estimate from internal summary statistics. Broadly, the CI p-values were less significant using external PRS, likely due to a combination of winner’s curse and the facts that external studies generally analyze datasets smaller than UKBB and do not perfectly match UKBB in terms of environment and genetic background. In addition to confirming our results, this analysis also suggests that CI can be reliably tested using internally-constructed PRS in the future. This can be essential for applications to less-studied traits, populations, or environmental contexts (Liu et al., 2018; Martin et al., 2019, 2017; Mefford et al., 2019; Mostafavi et al., 2019).

### Conservative step-down test adds confidence for Coordinated Interaction in UK Biobank

Despite our use of genetic PCs, cross-validation, and external summary statistics, we remain concerned about the possibility of confounding by residual population structure. As an additional robustness check, we re-analyzed these data with 10-fold lower p-value thresholds for the PRS. This is conservative because stricter p-value thresholds provide lower predictive power but almost certainly reduce the correlation between PRS and population structure.

We found broadly similar results, with most previous CI hits replicating (Supp Figure 2). Although several significant traits from the baseline analysis were no longer significant (internal height, SCV, LYM, TG, and external BMI and asthma), some of these traits retained support in their external/internal mirror image (e.g. BMI and asthma remain significant with internal PRS, and TG remains significant with external PRS). Moreover, the newly-insignificant traits were all near the threshold in the original analysis. Altogether, these results are consistent with moderate winner’s curse and reduced power from using a smaller set of SNPs as predictors, but the results do not suggest our EO test is severely inflated by population structure in the polygenic tail of small genetic effects.

### Tissue-specific coordination in UK Biobank

Having demonstrated the existence of CI for several traits, we now consider the possibility that interacting pathways are enriched in trait relevant tissues. Specifically, we test for tissue-specific enrichment of CI across the above 26 traits and 13 tissue-specific genomic annotations: 7 based on specifically-expressed genes (Adipocytes, Blood Cells, Brain, Hippocampus, Liver, Muscles, Pancreas) (Fehrmann et al., 2015; Pers et al., 2015), and 6 based on tissue-specific chromatin-marker patterns (Adipose, Brain, Hippocampus, Liver, Pancreas and Skeletal Muscle) (Roadmap Epigenomics Consortium et al., 2015; The ENCODE Project Consortium, 2012) as used in (Finucane et al., 2018).

For each tissue-trait pair, the tissue-specific EO test asks whether the coordination mediated through a specific tissue exceeds the genome-wide average. This test is conducted by first creating a tissue specific PRS on the even chromosomes (TPRS_*e*_) and testing it for interaction with the standard PRS on the odd chromosomes (PRS_*o*_). To ensure our test truly captures tissue-specific effects, rather than global effects that happen to be tagged by tissue-specific regions, we adjust for several covariates: the overall PRS_*e*_ ∗ PRS_*o*_, PRS_*e*_, PRS_*o*_, and TPRS_*e*_. Further, we adjust for the average TPRS across all tissues to test whether specific tissues are more trait-relevant than others and thus further reduce potential confounding by non-tissue-specific genomic annotations. We then repeat the process, flipping the roles of the even and odd PRS. Finally, we repeat the process with 50 random bifurcations (Methods). We again used a conservative Bonferroni threshold adjusting for both the number of splits and, now, also the number of tissues.

We found 7 instances of tissue-specific CI enrichment using internal PRS (Table 3). This includes liver enrichment for LDL, which is essentially a positive control given that many systems are involved in LDL and that the liver is a key regulator of LDL metabolism. Further confirming this result, liver-LDL was Bonferroni significant when using external PRS. The external PRS also suggestively implicate skeletal muscle and adipose, which are primary sinks for the triglycerides carried by LDL (Feingold and Grunfeld, 2000).

**Table 3.**
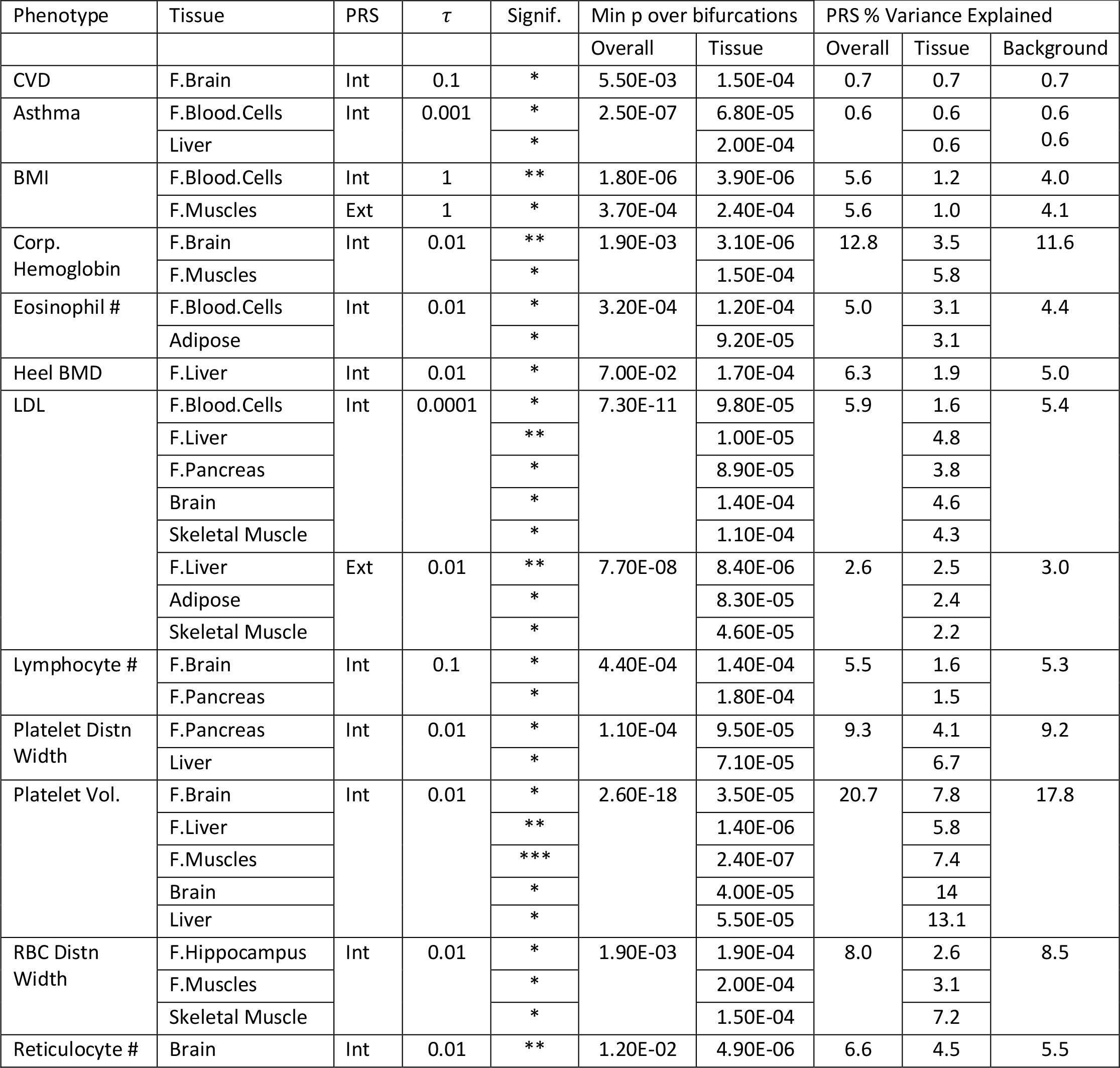
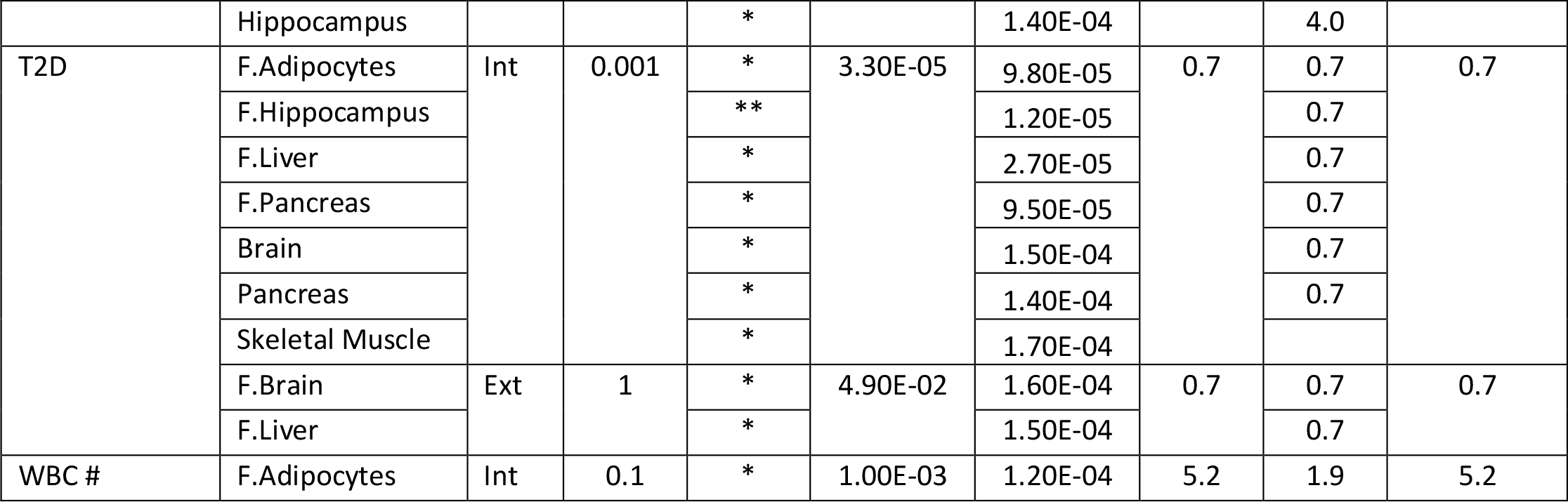
Tissue-specific CI in the UK Biobank. Trait-tissue pairs with significant tissue-specific Coordinated Interaction. All pairs where at least one chromosome split has p < 0.05/100 are shown (*); ** indicates p<.05/100/#tissues; and *** indicates p<.05/100/#tissues/#phenotypes. The prefix “F.” indicates that the tissue annotation is derived from Franke lab (Fehrmann et al., 2015; Pers et al., 2015). “Int” indicates that the PRS was calculated using the cross-validation method with UKBB data (Methods). “Ext” indicates that the PRS was calculated using GWAS summary statistics from external datasets (Methods). *τ* is the marginal SNP p-value threshold used to construct the PRS and is chosen to maximize cross-validated prediction accuracy.

Another biologically plausible set of tissue-trait pairs is for MPV, which also had the strongest ordinary CI of any trait. First, we find significant enrichment for liver, consistent with its known role as the main producer of the main regulator of platelet production, thrombopoietin (TPO) (Kaushansky, 2006). Second, we find significant CI enrichment for muscles, which also modestly produce TPO (Kaushansky, 2006); furthermore platelets are important in healing muscles. Third, we found suggestive CI with brain tissues—while the underlying biology is less obvious, there is evidence that TPO affects brain development (Ehrenreich et al., 2005). Additionally, recent complementary studies have found evidence that brain tissue plays a role in MPV (Barbeira et al., 2019).

We also found that T2D had significantly enriched CI in the hippocampus. This is interesting as the hippocampus is widely conjectured to play a role in memory and behavior (Anacker and Hen, 2017; Vikbladh et al., 2019), and the hippocampus is one of many brain tissues that has been linked to BMI (Cherbuin et al., 2015; Raji et al., 2010). Moreover, T2D is associated with physiological changes to the hippocampus (Ho et al., 2013), further supporting this CI enrichment and also emphasizing the complex interplay between tissues in determining disease and comorbidities. Broadly, these lines of evidence suggest the T2D-hippocampus CI is driven by a behavioral component of T2D. We caution, nonetheless, that hippocampus was the only specific brain region evaluated, and hence may primarily serve as a proxy for other brain regions; in particular, the hypothalamus has been robustly linked to metabolism and T2D (Smemo et al., 2014). The next three tissues prioritized for T2D—liver, pancreas, and adipose— are also obvious candidates for specific pathways to T2D: all are deeply involved in energy homeostasis, with the liver producing glucose, the pancreas producing insulin, and adipose serving as a primary sink for excess triglycerides (Stern et al., 2016). As additional support, we note that liver and hippocampus have suggestive evidence for CI when using the external PRS-based EO test.

Finally, another interesting, suggestive pair, blood cells-asthma, likely reflects the fact that we did not include any immune tissues and, hence, genes expressed in whole blood are the best available proxy for immune cells—which are known to play essential roles in asthma (Deckers et al., 2013).

In general, the tissue-specific EO test can improve power over the ordinary EO test when the correct tissue is identified, despite the fact that the TPRS almost always explains less trait variation than the overall PRS. As one example, red blood cell count is not remotely significant in the EO tests (p=1.50E-02), but its tissue-specific test with muscle is nearly significant (p=1.90E-04), even though the TPRS explains less than half the variance explained by the PRS (2.6% vs 8.0%, Table 3). Another example is mean corpuscular hemoglobin (MCH): although the EO test is insignificant (p=1.90E-03), the brain- and muscle-specific EO tests are significant (p=3.10E-06 and 1.50E-04) despite the fact that the TPRS again explain less than half of the variance explained by the overall PRS. Moreover, for MCH, the tissue with lower variance explained (brain, 3.5%) had stronger CI signal than the tissue with higher variance explained (muscle, 5.8%), again demonstrating the distinction between the signal sought by CI and the linear signal from percent variance explained. Overall, additional traits have significant CI after homing in on relevant tissues, and this happens despite the fact that the tissue-specific PRS have lower predictive power than ordinary PRS and, also, that we have conditioned on additional PRS-based covariates. This is further evidence that CI is truly enriched in specific tissues for some traits.

We chose to condition on the interaction between PRS_*o*_ and the across-tissue average TPRS_*e*_ when testing for tissue-specific CI. We found that the latter adjustment is important for uncovering trait-specific tissues because the unadjusted analysis prioritizes many tissues for a few traits with largest ordinary CI (Supplementary Figure 3): for example, the top 14 tissue-trait pairs contain 6 EA associations and 6 MPV associations. As tissue-specific annotations are partially confounded by other important genomic features, we hypothesize that conditioning on the average TPRS partially adjusts for these factors. In a related context, s-LDSC addresses this bias through the “baseline” genomic feature model (Finucane et al., 2018, 2015). More broadly, we caution that all standard caveats from genomic enrichment analyses apply—they are noisy, continually being improved, and liable to confounding from unadjusted genomic annotations, including both unmeasured tissues and more generic genomic features.

### Tissue-pair CI in UK Biobank

We next ask whether CI can be detected between pairs of tissues based on the hypothesis that pathways may be tissue specific. Rather than test the interaction between tissue-specific PRS*o* and global PRS*e*, we now test for interaction between tissue 1-specific PRS*o* and tissue 2-specific PRS*e*. For tissue-specific CI, we adjust for the same covariates as above, and now additionally adjust for PRS_*e*_ ∗ TPRS_*o*_ and TPRS_*e*_ ∗ PRS_*o*_. In particular, these tissue-x-tissue tests are statistically independent of the even-odd tests and the tissue-specific tests in the above sections under the null hypothesis for tissue-tissue interaction.

If tissue-x-tissue CI exists, it will likely cause tissue-specific CI, hence conditioning on tissue-trait pairs that are nominally significant will likely increase power. This reduction in testing dimension is particularly important for these tests because the number of tests multiplies by the number of evaluated tissues squared. We find several suggestive hits. This includes muscle-brain coordination for MPV, bolstering and elucidating the muscle-specific CI for this trait, which was by far the strongest signal in our tissue-specific EO test (Figure 3). LDL also had several suggestive results, including several interactions between liver and muscle, adipose, and whole blood. Similarly, T2D had several suggestive tissue interactions, including muscle, liver, and hippocampus.

**Figure 3.**
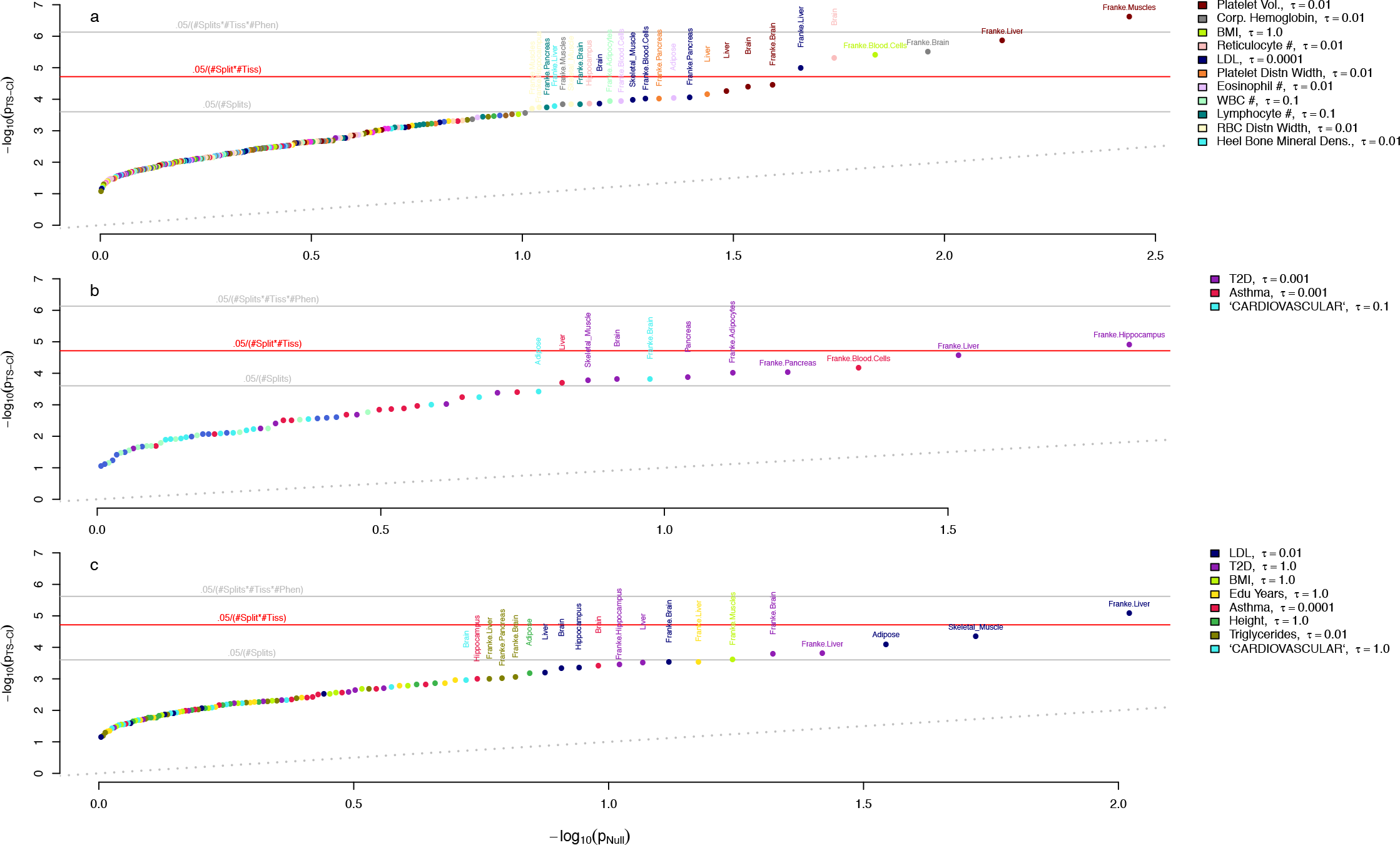
Tissue-specific Coordinated Interaction in the UK Biobank. Minimum p-value split for EO test using cross-validated in-sample PRS for 13 tissues and 21 quantitative traits (a) and 6 binary traits (b). (c) EO tests using external PRS for 8 of the quantitative and binary traits in (a) and (b) with external GWAS summary statistics available. In (a), the legend is only provided for the traits with highest mean –log10(p). *τ* is the p-value threshold used to construct the PRS and is chosen to maximize cross-validated prediction accuracy.

**Figure 4.**
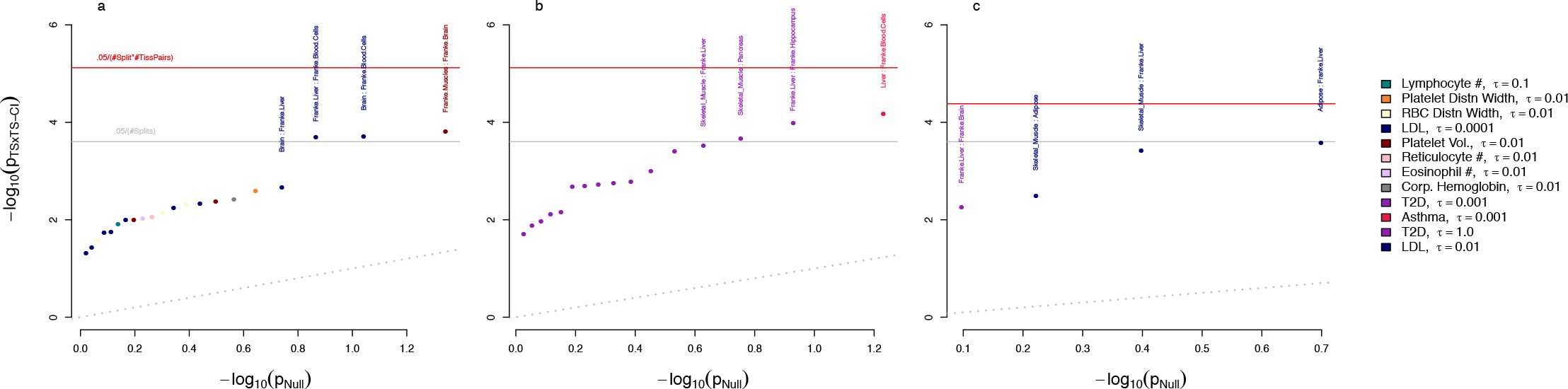
Candidate tissue-tissue CI in the UK Biobank. Points represent the minimum p-value for tissue-tissue CI over 100 splits of the chromosomes. Only tissue-trait pairs that are suggestively significant for tissue-specific CI are evaluated (i.e. p < 0.05/100 in Figure 3).

## Discussion

Epistatic interactions between genetic variants can have profound influences on phenotypic distributions and evolutionary landscapes (Phillips, 2008). While there are numerous examples of epistasis in model organisms and *in vitro* studies (Bloom et al., 2015; Brem et al., 2005; Corbett-Detig et al., 2013; Forsberg et al., 2017; Huang et al., 2012; Mackay, 2014), especially between pairs of genes, relatively few examples have been observed for complex polygenic traits in humans. In this work we propose a specific form of epistasis called Coordinated Interaction, in which many SNPs in one pathway jointly skew the effects of SNPs in other pathways. This model is inspired by recent theories of human genetic architecture (Boyle et al., 2017; Zuk et al., 2012). These pathways could represent distinct biological concepts, including different tissues (e.g. neurons and astrocytes in Alzheimer’s Disease) (Masters et al., 2015), different systems within a tissue (e.g. stress and immune response in Amyotrophic Lateral Sclerosis) (Phani et al., 2012), or unknown subtypes of a disease with partially distinct genetic risk factors (Dahl et al., 2019; Zuk et al., 2012). Importantly, coordination is fundamentally different from previous polygenic epistatic models: in particular, existing models assume a purely uncoordinated form of epistasis.

We next developed a test, called the *Even-Odd* test for Coordinated Interaction, based on testing the interaction between polygenic risk scores constructed from independent chromosomes. We showed via extensive analytical examination and empirical simulation that our test will detect Coordinated Interaction even though the pathways are unknown. We further show that our test is robust to (at least) moderate population structure and assortative mating. The sign of *γ* determines which direction CI skews the polygenic contribution, which is closely connected to the probability of observing extreme phenotypes in either direction. In the natural case where trait values are nonnegative, positive/negative epistasis corresponds to synergy/antagonism between main effects.

We observed evidence of Coordinated Interaction from the Even-Odd test for many phenotypes in the UKBB including blood, anthropometric, and disease phenotypes. We interpret this as evidence that Coordinated Interaction likely affects a broad range of phenotypes across many domains. To further investigate the Coordinated Interaction, we examined PRS restricted to genomic regions annotated to specific tissues and found biologically plausible results. Interestingly, for several phenotypes, the tissue-specific EO p-value increased in significance despite the fact that the total variance explained decreased. This suggests that tissue-specific annotations are enriched for true latent pathways in a trait-relevant way.

There are several limitations to this work and potential avenues to improving the Even-Odd test. First, more accurate PRS will improve the power of the test, either through more sophisticated models for PRS or through larger external reference datasets. Second, the EO test uses a random split of SNPs into even and odd PRS, which has low correlation with the true latent pathways and in turn reduces power. Using tissue-specific PRS is a step toward identifying which genomic regions harbor the greatest CI. Nonetheless, current variant-level annotations are imprecise, and other annotations may be more relevant depending on researcher interest and underlying trait biology. For example, annotations for cell-type specificity, gene ontology, inferred co-expression networks, or other functional categories related to methylation and chromatin state may be worth exploring. Third, the Bonferroni correction we use is overly conservative due to the strong correlation between splits (and tissues); some recent theoretical progress on testing multiple exchangeable hypotheses may enable power gains in the future (Rosset et al., 2018). Fourth, our random chromosome-splitting procedure itself has noise, and jointly testing all 210 pairs of chromosome-specific PRS interactions may add power; however, this is infeasible for modest data sets or for the extensions to test annotation-specific CI enrichment. Fifth, as with any test of interaction, phenotypic scale can impact results. However, we show that the sign and existence of CI is preserved under rescaling, either on average or under smooth transformation, and hence CI is (approximately) ‘essential’ (Sverdlov and Thompson, 2018). Sixth, we have not yet directly connected CI estimates to their impact on genetic architecture and heritability, though we do outline a path forward in the Supplementary Methods. Finally, as for all epistasis tests, the EO test only detects statistical CI, not biological CI. For example, a disease that has two subtypes with distinct, purely additive genetic bases will exhibit CI even though neither subtype has any epistasis at all. On the other side of the coin, though, CI can be a useful way to demonstrate the existence of unmodeled subtypes.

Going forward, several immediate extensions are interesting to explore. One could examine SNP x PRS interaction, or gene x PRS interaction as in TWAS (Gamazon et al., 2015; Gusev et al., 2016). This will add substantial power when the main effect of a highly-penetrant gene or SNP depends heavily on genetic background. Second, expanding to other traits and annotations could help refine the pathways involved and increasingly home in on causal mechanisms. For example, annotation of SNPs based on association with environmental factors, such as diet, exercise, smoking, or stress, can inform the nature of GxE interactions. Third, SNPs in genes known to influence specific pathways from in vitro molecular assays could be examined at the population scale in the full organismal context. For example, the genes in the Ribosomal Quality Control pathway are known to influences degradation of misfolded proteins (Brandman et al., 2012; Hickey et al., 2019), and this pathway may have important interactions with disease- or trait-specific coding variants segregating in the population. Finally, CI may have implications for medical interventions, which are generally pathway-specific. For example, if asthma exhibits strong positive CI between immune and lung pathways, intervening on either pathway will be sufficient to reduce disease burden; conversely, if LDL exhibits negative CI across liver- and BMI-driven pathways, it may be important to simultaneously address both pathways in order to control blood cholesterol.

## Methods

### Simulation description

Phenotypes and genotypes were generated under three phenotypic models (additive, uncoordinated interaction, and coordinated interaction) and three population models (random mating, assortative mating, and population structure) (Supplementary Methods). For each simulation, genotypes were generated for 2,000 individuals at 500 SNPs. Half of the individuals were randomly selected for the training set, in which the phenotype *y* was regressed on each genotype *g_j_* to calculate the estimated effect, 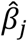, of each simulated SNP. The estimated effects of *g_j_* at each SNP were used as weights in the calculation of polygenic risk scores in the remaining 1,000 individuals 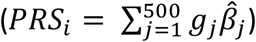. PRS were also calculated separately for even-indexed and odd-indexed SNPs.

### Model testing

To evaluate each population and phenotype model, linear regression was used to calculate the association between 1) the full PRS and phenotype, 2) the odd PRS and the even PRS, and 3) the interaction effect between the odd PRS and the even PRS on the phenotype. Under the assumption of random mating within a population, PRS calculated from unlinked SNPs (i.e. even/odd indexed) should be uncorrelated. To demonstrate this in our simulated populations, and to show that it does not hold under the scenario of assortative mating and population structure, the PRS calculated from even-indexed SNPs was regressed on PRS from odd-indexed SNPs. To test for models where the even-odd interaction between risk scores has a significant effect on phenotype, each phenotype was regressed on even PRS, odd PRS and an even by odd interaction term *y* ∼ *β*_0_ + *β*_1_*PRS_even_* + *β*_2_*PRS_odd_* + *β*_3_*PRS_even_ PRS_odd_*. The test was also performed conditional on the first 5 genetic PCs as covariates to evaluate this common approach to correct for population structure.

### UKBB data description

Data was obtained from the UK Biobank project (Bycroft et al., 2018). We used imputed genomic data version 3. We used the same phenotypic transformations as defined in (Finucane et al., 2018). Education level categories (data field #6138) were transformed to numerical values according to (Okbay et al. 2016 Nature) as follows: “College or University degree” = 20 years; “A levels/AS levels or equivalent” = 13 years; “O levels/GCSEs or equivalent” = 10 years; “CSEs or equivalent” = 10 years; “NVQ or HND or HNC or equivalent“ = 19 years; “Other professional qualifications eg: nursing, teaching” = 15 years; “None of the above” = 7 years; “Prefer not to answer” = missing. BMI was used as computed by UKBB (data field #21001). Heel bone mineral density (BMD) T-score was computed as the sum of the left and right Heel BMD T-score (data fields #4106 and 4125). Waist-hip Ratio was computed as the ratio of Waist circumference and Hip circumference (data fields #48 and 49). FEV1-FVC Ratio phenotype was computed as the ratio between data field ‘Forced vital capacity (FVC)’ (#3062) and ‘Forced expiratory volume in 1-second (FEV1)’ (#3063). T2D phenotype was based on data field ‘Diabetes diagnosed by doctor’ (#2443) where values as ’Prefer not to answer’ and ’Do not know’ were considered as missing data. Eczema allergy was based on data field ‘Blood clot, DVT, bronchitis, emphysema, asthma, rhinitis, eczema, allergy diagnosed by doctor’ (#6152) where the value ‘Hayfever, allergic rhinitis or eczema’ was considered to define cases, ’Prefer not to answer’ as missing, and the rest as controls. Asthma was based on the same data field, where only ‘Asthma’ values were considered as cases. High Cholesterol was determined based on samples with data code 1473 (‘high cholesterol’) in field ‘Non-cancer illness code, self-reported’ (#20002), all other samples were defined as controls. Cardiovascular cases were defined according to (Gazal et al., 2018) as samples having any of the following codes in field ‘Non-cancer illness code, self-reported’ (#20002): 1065 (hypertension); 1066 (heart/cardiac problem); 1067 (peripheral vascular disease); 1068 (venous thromboembolic disease); 1081 (stroke); 1082 (transient ischaemic attack (tia)); 1083 (subdural haemorrhage/haematoma); 1425 (cerebral aneurysm); 1473 (high cholesterol); 1493 (other venous/lymphatic disease).

### Sample selection

Sample QC: We filtered out samples who were not classified as ‘White British’ (data field 21000), or with discordance of genetic sex with declared sex (data fields 22001 and 31), or subjects who were identified as related to any other subject in the dataset as defined by UKBB (data field 22011). In total, 393076 subjects (180680 males, 212396 females) were included in further analysis.

Genetic QC: We filtered out variants with minor allele frequency (MAF) lower than 0.1%. In total, 16,525,182 variants were included in farther analysis. We clumped the variants based on MAF with r^2^ threshold of 0.2 and distance of 250kb. In total, 3,425,447 variants were selected after clumping. Clumping was performed using Plink1.9 (Chang et al., 2015). For other steps we used Plink v2.00a (Chang et al., 2015).

### Internal PRS

#### Cross validation by phenotype

For each phenotype, we split subjects into 10 folds. For each fold we estimated effect size of each variant using the remaining 9 folds, and we then use these effect sizes to construct a PRS for the held-out fold. Quantitative phenotypes were normalized for each fold separately by first removing outliers as samples outside 1.5 times the interquartile range above the upper quartile or below the lower quartile, and then performing quantile normalization. Although such normalization will inevitably shrink the non-Gaussianity induced by true CI signal, we view this as an important normalization step to remain conservative. Moreover, we prove smooth monotone transforms cannot create, remove, or change the sign of CI in the SOM.

PRS estimation consisted of two steps: (i) Effect size estimation; (ii) Compute PRS using selected variants. The first step is equivalent to performing a standard GWAS. For that purpose, we add as covariates the sex, age, center of assessment, genotyping batch, first 40 PCs as provided by UKBB, and BMI (except when BMI or T2D were the target phenotype). In the second step, PRS were computed for each fold for the 10% of samples that were not used to estimate the effect sizes. We computed PRS using a range of 11 P-values thresholds: 1.0, 0.1, 0.01, 0.05, 0.001, 0.0001, 1e-05, 1e-06, 1e-07, and 1e-08. Then for each phenotype, we chose the p-value threshold that maximized the percent variance explained in the held-out samples. We made an exception for asthma and used 0.0001, as the optimal choice (1e-05) gave zero weight to most chromosomes.

### External PRS

An alternative approach is to use effect sizes estimated from external datasets. This must be a cohort with similar ancestry in which the UKBB subjects were not included. We used external summary statistics included in LD Hub (Zheng et al., 2017) from the following studies: Height [GIANT consortium (Wood et al., 2014)]; LDL and Triglycerides [Global Lipids Genetics Consortium (Willer et al., 2013)]; BMI [GIANT consortium (Locke et al., 2015)]; educational attainment [Social Science Genetic Association Consortium (Okbay et al., 2016)]; T2D [DIAGRAM Consortium (Morris et al., 2012)]; Asthma [A GABRIEL Consortium (Moffatt et al., 2010)]; and CVD [CARDIoGRAMplusC4D Consortium (Nikpay et al., 2015)].

### Tissue-specific variants

Variants based on tissue-specific gene expression were defined as variants that fall within a 100kb of the genes that were classified as differentially expressed genes by (Fehrmann et al., 2015; Pers et al., 2015) similarly to (Finucane et al., 2018). Variants based on chromatin markers from the Roadmap project (Roadmap Epigenomics Consortium et al., 2015; The ENCODE Project Consortium, 2012). Any genomic location annotated with any of the DNase, H3K27ac, H3K36me3, H3K4me1, H3K4me3 and H3K9ac, was considered as ‘open chromatin’ region. To evaluate PRS based on tissue-specific variants, we used variants in the intersection of imputed variants from UKBB with MAF > 0.01% and variants that were classified as tissue-specific. We did not use the clumped set of variants, as the intersection of the two is small.

## Code Availability

Code to generate internal and external PRS; to perform CI and tissue-specific CI tests; and for simulations: https://github.com/nadavrap/CoordinatedInteractions/

## Acknowledgements

N.Z. is supported by NIH K25HL121295, U01HG009080, R01HG006399, R01CA227237, R03DE025665, and DoD W81XWH-16-2-0018. A.D. is supported by U01HG009080 and R01HG006399. B.S. is supported by R01CA227237, N.R. is supported by K25HL121295 and U01HG009080. S.J.S is supported by R01MH110928 and R01MH111662. This research has been conducted using the UK Biobank Resource under Application Number 30397.

---

**Supplementary Figure 1.**
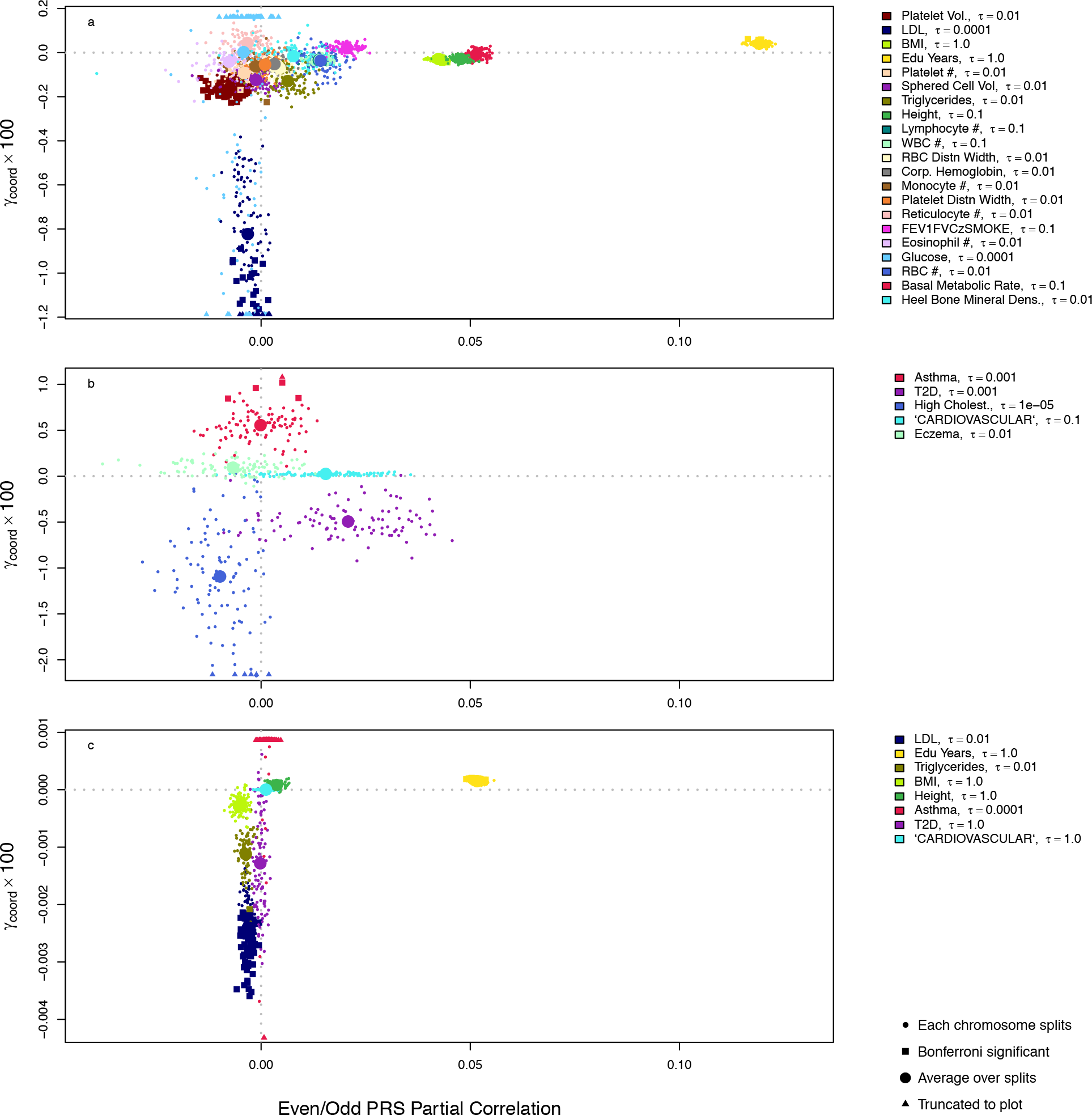
Coordinated Interaction estimates in the UK Biobank. Even-odd test statistics for CI using (a) cross-validated in-sample PRS for 21 quantitative traits (b) the same for 5 binary traits and (c) external PRS for 8 quantitative and binary traits. In (a), the legend is only provided for the traits with highest mean –log10(p). Each small point represents a single bifurcation of 22 autosomes. Bifurcations passing a strict Bonferroni threshold are highlighted as squares.

**Supplementary Figure 2.**
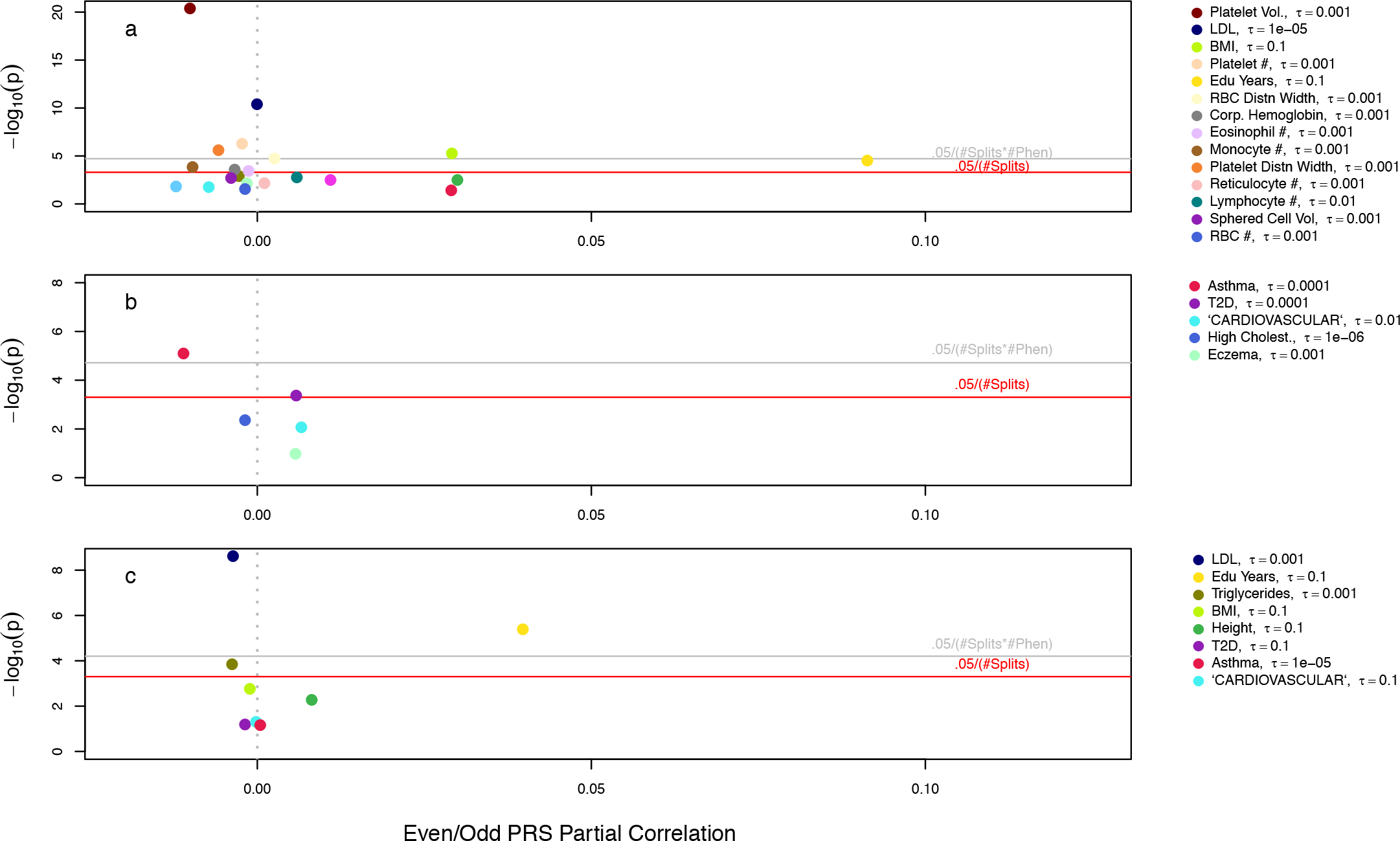
Coordinated Interaction in the UK Biobank at stricter thresholds. Even-odd tests for CI using (a) cross-validated in-sample PRS for 21 quantitative traits (b) the same for 5 binary traits and (c) external PRS for 8 quantitative and binary traits. In (a), the legend is only provided for the traits with highest mean –log10(p). Compared to main Figure 2, these analyses used 10-fold smaller p-value thresholds when constructing PRS.

**Supplementary Figure 3.**
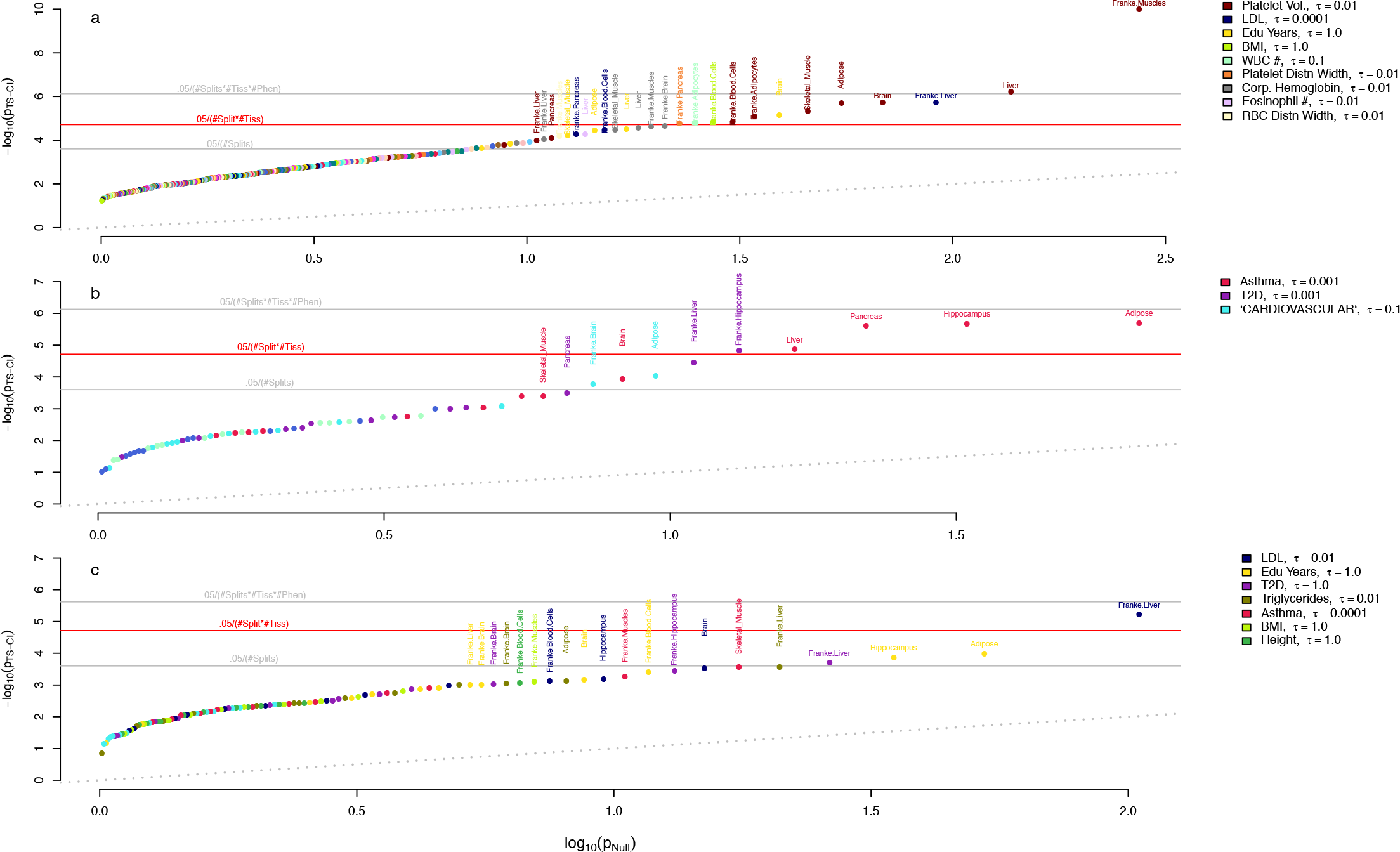
Tissue-specific Coordinated Interaction in the UK Biobank, unadjusted. Minimum p-value split for EO test using cross-validated in-sample PRS for 13 tissues and 21 quantitative traits (a) and 6 binary traits (b). (c) EO tests using external PRS for 8 of the quantitative and binary traits in (a) and (b) with external GWAS summary statistics available. In (a), the legend is only provided for the traits with highest mean –log10(p). *τ* is the p-value threshold used to construct the PRS and is chosen to maximize cross-validated prediction accuracy.

**Supplementary Table 1.** Polygenic simulations under additivity, uncoordinated interaction, or coordinated interaction assuming random mating, population structure, or assortative mating using correct SNPs in each group. Mean, standard deviation, and positive rate are shown for estimates of *θ_eo_* and *γ_eo_* for 10,000 simulations, where SNPs used to calculate PRS_e_ and PRS_*e*_ were the exact SNPs used to generate the phenotype. Positive rate (PR) is the proportion of significant test statistics at *α* = 0.05; sd is standard deviation.

| | Population Characteristics | | $\theta_{eo}$: $\text{PRS}_e \sim \text{PRS}_o$ | | | | $\gamma_{eo}$: $Y \sim \text{PRS}_e + \text{PRS}_o + \text{PRS}_e * \text{PRS}_o$ | | | |
| --- | --- | --- | --- | --- | --- | --- | --- | --- | --- | --- |
|  | Assortative Mating | Population Structure (FST = 0.1) | -PCs |  | +PCs |  | -PCs |  | +PCs |  |
| | | | $\chi^2$ (sd) | PR $\alpha=0.05$ | $\chi^2$ (sd) | PR $\alpha=0.05$ | $\chi^2$ (sd) | PR $\alpha=0.05$ | $\chi^2$ (sd) | PR $\alpha=0.05$ |
| Additive | 0 | 0 | 1.0 (1.4) | 0.0489 | -- | -- | 1.0 (1.4) | 0.0481 | -- | -- |
|  | 1 | 0 | 2.4 (2.8) | 0.222 | -- | -- | 1.0 (1.4) | 0.0517 | -- | -- |
|  | 0 | 1 | 6.7 (15.8) | 0.3231 | 1.0 (1.4) | 0.0487 | 1.0 (1.4) | 0.0475 | 1.0 (1.4) | 0.0493 |
| Uncoordinated Interaction | 0 | 0 | 1.0 (1.4) | 0.0471 | -- | -- | 1.0 (1.4) | 0.0535 | -- | -- |
|  | 1 | 0 | 2.4 (2.8) | 0.2152 | -- | -- | 1.0 (1.4) | 0.0503 | -- | -- |
|  | 0 | 1 | 6.8 (16.2) | 0.3257 | 1.0 (1.4) | 0.0472 | 1.0 (1.4) | 0.0518 | 1.0 (1.5) | 0.0549 |
| Coordinated Interaction | 0 | 0 | 1.0 (1.4) | 0.0484 | -- | -- | 185.2 (139.9) | 0.9437 | -- | -- |
|  | 1 | 0 | 3.7 (3.9) | 0.3646 | -- | -- | 291.1 (220.4) | 0.9527 | -- | -- |
|  | 0 | 1 | 7.9 (18.8) | 0.3343 | 1.0 (1.4) | 0.0498 | 167.7 (134.6) | 0.9335 | 180.1 (141.5) | 0.935 |

**Supplementary Table 2.**
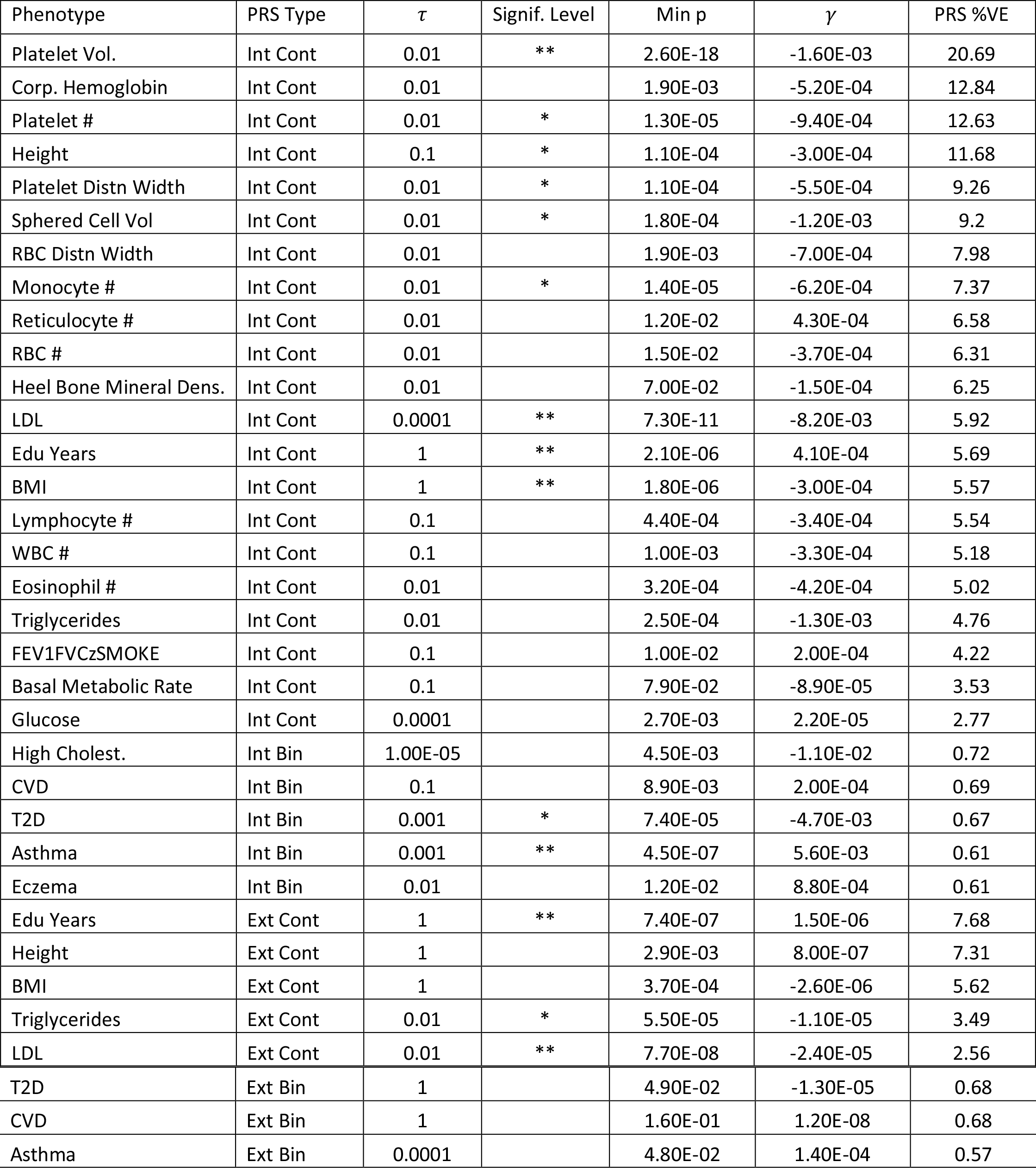
Even-odd test results for all traits in the UK Biobank. We summarize evidence for CI per trait using the minimum p-value over 100 random bifurcations, and provide the CI estimate (γ) for this top split. “Int” indicates that the PRS was calculated using the cross-validation method with UKBB data (Methods). “Ext” indicates that the PRS was calculated using GWAS summary statistics from external datasets (Methods). *τ* is the p-value threshold for the PRS and is chosen to optimize cross-validated prediction. PRS %VE describes the PRS variance explained.

