## Supplementary Text for "Coordinated Interaction: A model and test for globally signed epistasis in complex traits"

December 30, 2019

#### Outline

We first introduce our epistasis model and concept of *coordinated interaction* (CI). We then illustrate CI in the case of a nonnegative phenotype driven by two interacting SNPs (Section 1.1), where coordination is always perfect but may act either synergistically or antagonistically. Next, we study three SNPs with pairwise interactions (Section 1.2), which is sufficient to realize CI as a correlation across SNP pairs. In Section 2, we discuss how we estimate coordination by regressing on the interaction between even- and odd-chromosome PRS, as well as the key conditions underlying our Even/Odd approach. Section 3 lays out the interacting pathway model, where CI has a simple and interpretable form and the distribution of the Even/Odd estimator can be analytically derived. Finally, in Section 4, we describe several plausible biological models for coordination, as well as the behavior of our even/odd estimator Section 5 contains technical calculations.

### 1 Coordinated polygenic interaction

We assume a polygenic model with pairwise SNP interactions for a phenotype $y$ measured on $N$ samples. Let $G$ be the $N \times S$ genotype matrix of $S$ SNPs measured on the $N$ samples. We write the saturated pairwise interaction model as:

$$y_i \overset{\text{ind}}{\sim} \sum_{s=1}^{S} G_{is}\beta_s + \sum_{s \geq s'} G_{is}G_{is'}\Omega_{ss'} + \epsilon_i$$

where we define:

- $\beta \in \mathbb{R}^S$ is the vector of additive SNP effects.
- $\Omega$ is a matrix of pairwise SNP interaction effects. $\Omega_{ss'}$ gives the interaction effect between SNPs $s$ and $s'$. We assume $\Omega$ is lower triangular (WLOG).
- We assume that $\epsilon$ has i.i.d. Gaussian entries with mean zero.
- We assume columns of $G$ are scaled to mean 0, variance 1 (WLOG in the sense that $\beta$ and $\Omega$ can be rescaled concomitantly).
- We have excluded fixed effect covariates for notational convenience.
- We also assume columns of $G$ are independent. Allowing LD would complicate calculations and estimation details but, we suspect, would not fundamentally change the situation.

The model can be written more succinctly by using $*$, the column-wise Khatri-Rao product. Each column of the Khatri-Rao matrix product $A * B$ is equal to the element-wise product between a column of $A$ and a column of $B$. Although apparently complicated, the Khatri-Rao product is a basic component of linear regression with interactions, e.g. $A * B$ is exactly what `R` creates with `model.matrix(~-1+A:B)`. Using $*$, the pairwise interaction model is:

$$y \sim G\beta + (G * G)\,\omega + \epsilon \tag{1}$$

This defines $\omega := \text{vec}(\Omega) \in \mathbb{R}^{S^2}$, where $\text{vec}(\cdot)$ concatenates the columns of a matrix into a vector.

Previous analyses of such pairwise SNP interaction models have focused on two distinct, complementary strategies. First, individual pairwise SNP effects can be tested by fitting individual SNP pairs (or higher-order tuples) as fixed effects [1]. Second, random effect models can be used to aggregate all SNP effects and SNP-SNP interaction effects under the assumption that these effects are all independent [2–10]. The former approach is particularly attractive for SNPs of large effect or candidate SNPs, but genome-wide screens can also be useful even in some polygenic contexts [11]. The latter approach generally has high power for complex traits, but sacrifices resolution on the specific causal SNPs and nature of the interactions. Both approaches have been established for decades.

Several recent approaches have worked to bridge this gap. A particularly simple and elegant approach is to test the interaction between a single SNP and a polygenic random effect [12]–on one hand, this provides the resolution of SNP-level tests, and, on the other, borrows strength across the entire genome. A related concept is to test for interaction between SNPs and/or PRS and rich, partially unknown environments, which has been facilitated either by assuming that the latent environment is univariate [13]–hence admits simple analytic marginalization–or is multivariate with prior covariance given by known proxy environmental measures [14].

We introduce a new type of polygenic interaction called coordination, defined (roughly) as:

$$\gamma(\Omega, \beta) = \mathrm{Cor}_{s \neq s'} \left( \Omega_{ss'}, \beta_s \beta_{s'} \right) \tag{2}$$

Intuitively, coordination measures whether the interactions line up with the product of main effects. When $\gamma > 0$, the interactions in $\Omega$ work in concert to amplify positive SNP effects; for $\gamma < 0$, however, $\Omega$ dampens positive SNP effects. We call the coordination *perfect* if $\gamma \in \{-1, 1\}$. The definition of coordination excludes the diagonal terms $\Omega_{ss}$ because we want to capture interactions between different SNPs, not nonlinear single-SNP effects like dominance.

We next illustrate the notion of coordination in the simple cases of two and three SNPs. With two SNPs, we can show how coordination measures the synergy/dampening of main SNP effects. With three SNPs, we can show how polygenic coordination measures the average of the synergy/dampening across all causal SNP pairs. We assume nonnegative phenotypes in these stylistic sections so that synergy/antagonism are equivalent to positive/negative epistasis [15]. Although coordination generally measures the latter, the former can be more intuitive.

In Section 4, we provide several plausible polygenic models and show how they induce CI:

- Master transcription factors
- Polygenic buffers
- Structured random $\Omega$
- GxE with G-E dependence

#### 1.1 Pairwise interaction between two SNPs

In these two subsections, we write $\Omega_{ss'} = \beta_s \beta_{s'} \tau_{ss'}$; this notation is useful and comes without loss of generality except in aberrant situation where a SNP has zero marginal effect and nonzero interaction effects.

The exact definition of *coordination* is:

$$\gamma(\Omega, \beta) := \frac{\sum_{s>s'} \Omega_{ss'} \beta_s \beta_{s'}}{\sqrt{\sum_{s>s'} \Omega_{ss'}^2} \sqrt{\sum_{s>s'} \beta_s^2 \beta_{s'}^2}} \tag{3}$$

Although (3) is our exact definition of coordination, we usually think about the correlation-based approximation in (2) because it is simple, interpretable and accurate for the polygenic models.

Our example imagines two haploid SNPs a/A and b/B with main effects $\beta_A$ and $\beta_B$. To ease interpretation, assume that the SNPs are coded such that $\beta_A, \beta_B > 0$ and that the mean phenotype for the ab genotype is 0. We parameterize the pairwise interaction through $\tau_{AB}$, which describes the gap between the true interaction model and the linear prediction for the mean AB phenotype (Table 1). In other words, the mean phenotype for samples with the AB genotype is $\beta_A + \beta_B$ under the linear model, but in reality the mean phenotype for AB samples is $\beta_A + \beta_B + \beta_A \beta_B \tau_{AB}$. Hence, $\tau_{AB} < 0$ means the genotypes act antagonistically, with the total effect of $A$ and $B$ being less that the sum of its parts. On the other hand, $\tau_{AB} > 0$ means the SNPs act synergistically. In the former case, adding marginal SNP effects overestimates the true genetic variance explained; in the latter case, adding marginal SNP effects fails to capture all the (broad-sense) heritability.

For a single pair of SNPs, the coordination is trivially perfect by the definition in (3) (unless $\tau_{AB}$ = 0, in which case there is no interaction to coordinate). This emphasizes the difference between coordination and generic interaction: rather than measure the existence or size of interactions, coordination assumes interactions exist and measures their alignment with main effects.

| | a | A |
|---|---|---|
| b | 0 | $\beta_A$ |
| B | $\beta_B$ | $\beta_A + \beta_B + \beta_A\beta_B\tau_{AB}$ |

$$\implies \gamma = \frac{\beta_A^2\beta_B^2\tau_{AB}}{\sqrt{\beta_A^2\beta_B^2\tau_{AB}^2}\sqrt{\beta_A^2\beta_B^2}} = \text{sign}(\tau_{AB})$$

Table 1: Mean phenotype values with two interacting, haploid SNPs (a/A and b/B) with baseline effects $\beta_A, \beta_B > 0$. When $\tau < 0$, the AB genotype is buffered relative to the additive model; when $\tau > 0$, the genetic effect is super-linear. $\gamma$ is formally zero when the linear model holds.

#### 1.2 Pairwise interaction between three SNPs

In this case of two SNPs, there is only one interaction effect, hence the coordination is trivially perfect: all SNP pairs (i.e. the only SNP pair) have either inflated- or deflated-effects relative to linearity, hence the coordination is perfect, with sign determined by $\tau$. More generally, if there are many SNPs, $\gamma$ is like a weighted average over the $\gamma$ values for each SNP pair. Intuitively, $\gamma$ estimates whether positive SNP effects are, on balance, inflated or deflated (relative to additive) in combination with another SNP.

We illustrate this in the case of three SNPs in Table 2. The first result is that $\gamma$ decomposes into a weighted average of the $\tau$ for each SNP pair (the † approximation in Table 2 below). This allows us to extend our intuition from the two-/three-SNP cases to the more general polygenic models in later sections.

The second result is that $\gamma^2$ roughly describes the proportion of genetic interaction that is coordinated with main effects:

$$\gamma^2 \approx \frac{\bar{\tau}^2}{\bar{\tau}^2 + \mathbb{V}(\tau)} \tag{4}$$

This is formalized and generalized to polygenic models in Section 5.1.

| | a | A |
|---|---|---|
| b | 0 | $\beta_A$ |
| B | $\beta_B$ | $\beta_A + \beta_B + \beta_A\beta_B\tau_{AB}$ |

| | a | A |
|---|---|---|
| c | 0 | $\beta_A$ |
| C | $\beta_C$ | $\beta_A + \beta_C + \beta_A\beta_C\tau_{AC}$ |

| | c | C |
|---|---|---|
| b | 0 | $\beta_C$ |
| B | $\beta_B$ | $\beta_B + \beta_C + \beta_B\beta_C\tau_{BC}$ |

Define $w_{ll'} := \frac{\beta_l^2\beta_{l'}^2}{\sqrt{\sum_{s>s'}\tau_{ss'}^2\beta_s^2\beta_{s'}^2}\sqrt{\sum_{s>s'}\beta_s^2\beta_{s'}^2}}$

$$\begin{aligned}\implies \gamma &= w_{AB}\tau_{AB} + w_{BC}\tau_{BC} + w_{AC}\tau_{AC} \\ &= \sum_{s>s'} w_{ss'}\tau_{ss'} \\ &\overset{\dagger}{\approx} \frac{\bar{\tau}}{\sqrt{\bar{\tau}^2 + \mathbb{V}(\tau)}} \\ \implies \gamma^2 &\approx \frac{\bar{\tau}^2}{\bar{\tau}^2 + \mathbb{V}(\tau)}\end{aligned}$$

Table 2: The equalities are general and emphasize that $\gamma$ is a type of weighted genome-wide average across all SNP pairs. The approximation works well for polygenic models, see Section 5.1.

#### 2 The even/odd estimator for coordination

Clearly, if we knew $\Omega$ and $\beta$ we could just calculate the coordination $\gamma$ using equation (3). More generally, we could evaluate $\gamma$ using estimates of $\Omega$ and $\beta$.

However, we are primarily interested in the domain where $S \gg N$ and $\beta$ and $\Omega$ are dense, a setting where accurate effect estimates are hopeless. In the simpler context of GREML, where $\Omega$ is omitted, this high-dimensionality has led many others to consider random effect models to estimate $\|\beta\|$ directly, without first estimating $\beta$. GREML works well for polygenic signals because (a) there are more causal SNPs than samples and (b) $\|\beta\|$ can be accurately estimated even when the assumed parametric prior for $\beta$ is relatively badly misspecified [16, 17].

Redolent of GREML, we do not attempt to fit all SNP effects and pairwise SNP-SNP interactions. Instead, we average these signals over large genomic regions. For sufficiently polygenic traits, this averaging increases power without incurring bias. Intuitively, both approaches succeed when nonzero elements of $\beta$ (and $\Omega$, for CI) are spread uniformly across the genome, which happens with high probability for polygenic traits. We formalize this under a model where phenotypes are driven by interactions between additively-heritable pathways in Section 3.

Specifically, we are interested in the regression coefficients derived from fitting the following (misspecified) phenotypic model with ordinary linear regression, symbolically represented by:

$$y \sim \alpha_A P_A + \alpha_B P_B + \lambda P_A * P_B \tag{5}$$

$$P_{iA} := \sum_{s \in A} G_{is}\beta_s; \quad P_{iB} := \sum_{s \in B} G_{is}\beta_s \tag{6}$$

This defines $P_A$ and $P_B$ as the risk score derived from the SNP sets $A$ and $B$. For example, we are primarily motivated by the case where $A =$ SNPs on odd chromosomes, and $B =$ even chromosomes.

We propose to estimate the coordination $\gamma$ by rescaling the OLS estimate of $\lambda$:

$$\hat{\gamma}_{AB} := \sqrt{\frac{h_A^2 h_B^2}{h_{AB}^2}} \hat{\lambda}_{OLS} \tag{7}$$

$$h_A^2 := \|\beta_A\|^2; \quad h_B^2 := \|\beta_B\|^2; \quad h_{AB}^2 := \|\Omega_{AB}\|^2$$

Although this scaling is essential for unbiasedly estimating $\gamma$, it has no effect at all on testing whether $\gamma$ is zero or the estimated sign of $\gamma$. For this reason, we focus on $\hat{\lambda}$ in the main text (though we call it $\hat{\gamma}$ for simplicity), and we only evaluate CI through its sign and $p$-value. In the future, it may be useful to scale $\lambda_{OLS}$ using estimates of $h_A^2$, $h_B^2$ and $h_{AB}^2$ derived from models like [2–10], which may be helpful for quantifying the impact of CI on genetic architecture.

Our main theoretical result (Proposition 1) shows that this estimator is unbiased and consistent:

$$\hat{\gamma}_{AB} = \gamma_{AB} + \mathcal{N}\left(0, \frac{1}{N h_{AB}^2}\right)$$

where the true $A/B$ odd coordination $\gamma_{AB}$ is defined as:

$$\gamma_{AB} = \frac{\beta_A^T \Omega_{AB} \beta_B}{\sqrt{h_A^2 h_B^2 h_{AB}^2}}$$

Intuitively, $N h_{AB}^2$ is the effective sample size for coordination estimation.

This result provides the link between our estimated even/odd CI and the true even/odd CI, i.e. $\hat{\gamma}_{AB} \approx \gamma_{AB}$. In the next section, we will ask which models yield $\gamma_{AB} \approx \gamma$. Together, this will show that $\hat{\gamma}_{AB}$ is a reasonable estimator of the coordination $\gamma$.

We assumed the true $\beta$ are used in constructing these risk scores. We expect that errors in estimating $\beta$ will attenuate $\gamma$ estimates and power but will not induce bias or false positives, similar to genotyping or phenotyping errors in GREML [17]. Tracking the effects of errors in $\beta$ seems labor intensive but straightforward. Also, we assume that the pathways are independent, i.e. $A$ and $B$ are disjoint. We lift this in Section 3.3.

The interaction between polygenic scores in model (5) coincides with the fully saturated interaction model in (1) when there exists some $\lambda \in \mathbb{R}$ such that:

$$\Omega = \begin{pmatrix} 0 & \Omega_{(21)} \\ \Omega_{(12)} & 0 \end{pmatrix} = \lambda \begin{pmatrix} 0 & \beta_{(2)}\beta_{(1)}^T \\ \beta_{(1)}\beta_{(2)}^T & 0 \end{pmatrix} \tag{8}$$

We call this perfect even/odd coordination. This implies perfect (general) coordination, and essentially adds the further constraint that all coordination flows through the specific even/odd partition.

### 3 Interacting pathways model

Suppose the trait is defined by pairwise interactions between $K$ discrete system-/pathway-specific contributions. In this section, we derive the coordination, $\gamma$, and show the even/odd estimator, $\hat{\gamma}_{AB}$, is accurate for polygenic models. We then discuss several biological mechanisms that yield such interacting pathway models in Section 4.

We formalize the interacting pathways model as:

$$y = \sum_k z_k + \sum_{k \leq k'} (z_k \circ z_{k'})\, \alpha_{kk'} + \epsilon \tag{9}$$

$$z_k = \sum_{s \in S_k} G_{,s}\beta_s$$

where $z_k$ are the $K$ discrete pathway main effects driven by the SNPs in the the set $S_k$. Note that $z$ reflect causal biological pathways, while the $A/B$ risk scores we construct in $P$ are based on chosen, potentially arbitrary SNP sets.

Note that this interacting pathways model in (9) is an instance of the full polygenic pairwise interaction model in (1), with $\omega_{ss'} = \beta_s\beta_{s'}\alpha_{k(s)k(s')}$, where $k(s)$ indicates the pathway to which SNP $s$ contributes.

##### 3.1 $\gamma$ under interacting pathways

In the interacting pathways model, the coordination is:

$$
\begin{aligned}
\gamma &:= \frac{\sum_{s<s'} \beta_s \beta_{s'} \omega_{ss'}}{\sqrt{\left(\sum_{s<s'} \beta_s^2 \beta_{s'}^2\right)\left(\sum_{s<s'} \omega_{ss'}^2\right)}} \\
&= \frac{\sum_{s<s'} \beta_s^2 \beta_{s'}^2 \alpha_{k(s)k(s')}}{\sqrt{\left(\sum_{s<s'} \beta_s^2 \beta_{s'}^2\right)\left(\sum_{s<s'} \beta_s^2 \beta_{s'}^2 \alpha_{k(s)k(s')}^2\right)}} \\
&= \frac{\sum_{k,k'} h_k^2 h_{k'}^2 \alpha_{kk'} - \sum_s \beta_s^4 \alpha_{k(s)k(s)}}{\sqrt{\left(\sum_{k,k'} h_k^2 h_{k'}^2 - \sum_k \sum_{s\in k} \beta_s^4\right)\left(\sum_{k,k'} h_k^2 h_{k'}^2 \alpha_{kk'}^2 - \sum_s \beta_s^4 \alpha_{k(s)k(s)}^2\right)}} \\
&=: \frac{\sum_{k,k'} h_k^2 h_{k'}^2 \alpha_{kk'} - s_1}{\sqrt{\left(h_\beta^4 - s_2\right)\left(h_\omega^2 - s_3\right)}} \\
&\approx \frac{\sum_{k,k'} h_k^2 h_{k'}^2 \alpha_{kk'}}{h_\beta^2 \sqrt{h_\omega^2}}
\end{aligned}
$$

$$
h_k^2 := \sum_{s\in k} \beta_s^2; \quad h_\beta^2 := \sum_s \beta_s^2 = \sum_k h_k^2; \quad h_\omega^2 := \sum_{k,k'} h_k^2 h_{k'}^2 \alpha_{kk'}^2 \approx 2 \sum_{s<s'} \omega_{ss'}^2
$$

A few facts:

- $\gamma$ is invariant to jointly multiplying all $h_k^2$ or all $\alpha_{kk'}$ by any positive number.
- $\gamma \neq 0$ implies that at least one heritable, interacting pathway exists–$\exists\, i,j$ s.t. $h_i^2, h_j^2, \alpha_{ij} \neq 0$.
- For $K = 2$, with no intra-pathway nonlinearity (i.e. $\alpha_{11} = \alpha_{22} = 0$),

$$
\gamma \approx \frac{h_1^2 h_2^2 \alpha_{12}}{(h_1^2 + h_2^2)\sqrt{h_1^2 h_2^2 \alpha_{12}^2}} = \operatorname{sign}(\alpha_{12}) \frac{\sqrt{h_1^2 h_2^2}}{h_1^2 + h_2^2} \tag{10}
$$

  which is maximized in absolute value when $h_1^2 = h_2^2$.

- Assume all pathways interact equally in that $\alpha_{ij} = aI\{j \leq i\}$. Then:

$$
\gamma = \operatorname{sign}(a) \sqrt{\frac{\sum_{k\leq k'} h_k^2 h_{k'}^2}{\sum_{k,k'} h_k^2 h_{k'}^2}} = \operatorname{sign}(a)\sqrt{1/2}\sqrt{1 + \frac{\sum_k h_k^4}{\left(\sum_k h_k^2\right)^2}}
$$

  this attains its maximum, $|\gamma| = 1$, when $h_k^2 \neq 0$ for exactly one $k$. If instead $\alpha_{ij} = aI\{j < i\}$, the coordination is:

$$
\gamma = \operatorname{sign}(a) \sqrt{\frac{\sum_{k<k'} h_k^2 h_{k'}^2}{\sum_{k,k'} h_k^2 h_{k'}^2}} = \operatorname{sign}(a)\sqrt{1/2}\sqrt{1 - \frac{\sum_k h_k^4}{\left(\sum_k h_k^2\right)^2}}
$$

  which has opposite properties: $|\gamma|$ is maximized at $\sqrt{\frac{K-1}{2K}}$ when all $h_k^2$ are equal, and $\gamma = 0$ when all $h_k^2 = 0$ except for one $k$.

- Assume $K$ is large and $h^2$ and $\alpha$ are independent. Then:

$$\begin{aligned}\gamma &\approx \frac{\sum_{k,k'} h_k^2 h_{k'}^2 \alpha_{kk'}}{h_\beta^2 \sqrt{h_\omega^2}} \\ &\approx \frac{K^2 \left(\frac{1}{K^2}\sum_{k,k'} h_k^2 h_{k'}^2\right)\left(\frac{1}{K^2}\sum_{k,k'} \alpha_{kk'}\right)}{h_\beta^2 \sqrt{K^2 \left(\frac{1}{K^2}\sum_{k,k'} h_k^2 h_{k'}^2\right)\left(\frac{1}{K^2}\sum_{k,k'} \alpha_{kk'}^2\right)}} \\ &= \frac{h_\beta^4 \bar{\alpha}}{h_\beta^2 \sqrt{h_\beta^4 \left(\bar{\alpha}^2 + \mathbb{V}(\alpha)\right)}} \\ &= \frac{\bar{\alpha}}{\sqrt{\bar{\alpha}^2 + \mathbb{V}(\alpha)}}\end{aligned}$$

  In this case, $\gamma$ is the average level of positive/negative epistasis across pathways, shrunk toward zero to penalize variance between pairwise pathway-level interactions.

- We have loosely assumed that the residual terms $s_1$, $s_2$, and $s_3$ are approximately zero. This will hold when the heritability is smoothly spread across many causal SNPs because these terms each sum over only $L$ SNP pairs rather than $L^2$. This is particularly easy to see in the limit that $\beta_s^2 = b$ for all SNPs $s$. Taking $s_1$ as an example, and defining $L_k$ as the number of SNPs contributing to pathway $k$:

$$\left|\frac{s_1}{\sum_{s,s'} \beta_s^2 \beta_{s'}^2 \alpha_{k(s)k(s')}}\right| = \left|\frac{2\sum_s b^2 \alpha_{k(s)k(s)}}{\sum_{s,s'} b^2 \alpha_{k(s)k(s')}}\right| = 2\left|\frac{\sum_k L_k \alpha_{kk}}{\sum_{k,k'} L_k L_{k'} \alpha_{kk'}}\right| \leq 2|\alpha_{(1)(1)}|/L_{(1)}$$

  where the inequality is over choices of $\alpha$ and (1) indicates the value of $k$ that minimizes $L_k$. So, as long as we consider the limit $(N, L_j) \to \infty$, then $s_1, s_2, s_3 \to 0$.

- In the large-$K$ limit and assuming $\alpha$ has mean 0, the coordination approximately becomes a type of pathway-level coordination: letting $h := \left(\sqrt{h_1^2}, \ldots, \sqrt{h_K^2}\right)$,

$$\gamma = \mathrm{Corr}\left(h \otimes h, (h \otimes h) \circ \mathrm{vec}(\alpha)\right)$$

  If the pathway interaction strengths ($\alpha$) are not coordinated with the product of main pathway effects ($h \otimes h$), then the coordination vanishes. In other words, coordination only persists in the large $K$ limit when the pathways themselves are coordinated. Biologically, this is not expected to be a particularly interesting limit, which motivates our focus on traits driven by interactions between a small number of pathways.

#### 3.2 $\gamma_{AB}$ under interacting pathways

Above, we derived the behavior of the true, global coordination, $\gamma$. Here, we analyze $\gamma_{AB}$, the oracle estimator of $\gamma$ we could create if we perfectly knew $\beta_A$, $\beta_B$, and $\Omega_{AB}$. Starting from its definition,

$$
\begin{aligned}
\gamma_{AB} &:= \frac{\beta_A^T \Omega_{AB} \beta_B}{\sqrt{h_A^2 h_B^2 h_{AB}^2}} \\
&= \frac{\sum_{s\in A, s'\in B} \beta_s (\beta_s \beta_{s'} \alpha_{k(s)k(s')}) \beta_{s'}}{\sqrt{h_A^2 h_B^2}\sqrt{\sum_{s\in A, s'\in B} \left(\alpha_{k(s)k(s')} \beta_s \beta_{s'}\right)^2}} \\
&= \frac{\sum_{k,k'} \alpha_{kk'} \sum_{s\in A\cap k, s'\in B\cap k'} \beta_s^2 \beta_{s'}^2}{\sqrt{h_A^2 h_B^2}\sqrt{\sum_{k,k'} \alpha_{kk'}^2 \sum_{s\in A\cap k, s'\in B\cap k'} \beta_s^2 \beta_{s'}^2}} \\
&= \frac{\sum_{k,k'} \alpha_{kk'} h_{A,k}^2 h_{B,k'}^2}{\sqrt{h_A^2 h_B^2}\sqrt{\sum_{k,k'} \alpha_{kk'}^2 h_{A,k}^2 h_{B,k'}^2}}
\end{aligned}
$$

We have defined $h_{A,k}^2 = \sum_{s\in A\cap S_k} \beta_s^2$ as the pathway-$k$ heritability that is captured by the SNPs in group $A$ (and likewise for $h_{B,k}^2$).

If only one pathway pair is active–$\alpha_{12} \neq 0$, say–then:

$$
\gamma_{AB} = \text{sign}(\alpha_{12}) \sqrt{\frac{h_{A,1}^2}{\sum_k h_{A,k}^2}} \sqrt{\frac{h_{B,2}^2}{\sum_k h_{B,k}^2}} \tag{11}
$$

showing $\gamma_{AB}^2$ measures the fraction heritability in group $A$ ($B$) that tags pathway 1 (2). Hence $\gamma_{AB}$ increases either when (a) superfluous main effects are excluded, so that $h_{A,k}^2$ ($h_{B,k'}^2$) decreases for $k \neq 1$ ($k' \neq 2$) or (b) meaningful main effects are added, so that $h_{A,1}^2$ or $h_{B,2}^2$ increases.

We note that for non-perfectly-polygenic models, our assumption that $\alpha_{21} = 0$ is not trivial, as $A$ and $B$ may differ meaningfully. In this case, $\Omega$ must be symmetrized (i.e. we must set $\alpha_{21} = \alpha_{12}$ and then divide by 2). In this case,

$$
\gamma_{AB}^2 = \frac{1}{2} \frac{h_{A,1}^2 h_{B,2}^2 + h_{A,2}^2 h_{B,1}^2}{h_A^2 h_B^2} \tag{12}
$$

The is useful for the master transcription factor model in 4.1, because the master TF is assumed to derive all its heritability from a small, contiguous genomic region that must live entirely in either $A$ or $B$. It is also clear that this exactly reduces to (11) under perfect polygenicity, as then $h_{A,2}^2 h_{B,1}^2 = h_{B,2}^2 h_{A,1}^2$.

Compared to $\gamma$ in (10), $\gamma_{AB}$ replaces the proportion of heritability tagged by group 1 ($h_1^2/h_\beta^2$) with the proportion of $A$-heritability tagged by group 1 ($h_{A,1}^2/h_A^2$); the $B$ group is analogous. In other words, $\gamma_{AB}/\gamma$ measures the enrichment of coordination between $A$ and $B$ SNPs compared to the average coordination across all SNP pairs. Intuitively, most choices for $A$ and $B$ will give $\gamma_{AB}$ that reasonably approximates $\gamma$.

In the special case that the $A/B$ split is purely random and the system is highly polygenic, then $h_{A,k}^2 \approx \mathbb{E}_{A/B\ split}\left(h_{A,k}^2\right) = \frac{|A|}{S} h_k^2$ (and $h_{B,k}^2 \approx \frac{|B|}{S} h_k^2$), which means that the $AB$ estimator is approximately unbiased: $\mathbb{E}_{A/B\ split}(\gamma_{AB}) \approx \gamma$. This informally establishes that random SNP partitions yield accurate estimates of polygenic coordination.

However, when $A$ and $B$ are chosen adversarially, arbitrarily large bias can arise (as in GREML, [17]). For example, when $A$ perfectly captures pathway 1 heritability ($h_{A,1}^2 = h_1^2$) and only pathway

1 heritability ($h^2_{A,1} = h^2_A$), and analogous for $B$ perfectly coinciding with pathway 2, then $\gamma^2_{AB} = 1$ regardless the overall polygenic coordination.

More generally, increasing the concentration of $A$ around pathway 1 (resp. $B$ around 2) increases the $AB$ coordination. In this sense, we can use the variation of $\gamma_{AB}$ as a function of $A$ and $B$ to map the latent interacting pathways. This is the intuition supporting our tests for enrichment of CI in trait-relevant tissues, but can be generalized in many ways. For example, in the limit where $|A| \to 1$, a SNP-level model is obtained, giving a coordination variant of the SNP-by-Polygenic interaction test in [12].

##### 3.3 Incorporating genetically correlated pathways

Now, we no longer require that the pathways $z_k$ are each determined by non-overlapping sets of SNPs. Instead, we just assume that each pathway follows an additive model, with pathway $k$ determined by the effects $\beta^{(k)}$:

$$z_k = G\beta^{(k)}$$

In the specific case of $K = 2$, assuming no intra-pathway non-linearity ($\alpha_{11} = \alpha_{22} = 0$), the coordination is now:

$$\begin{aligned}
\gamma &:= \frac{\sum_{s<s'}(\sum_k \beta_s^{(k)})(\sum_k \beta_{s'}^{(k)})\left(\alpha_{12}\beta_s^{(1)}\beta_{s'}^{(2)}\right)}{\sqrt{\left(\sum_{s<s'}(\sum_k \beta_s^{(k)})^2(\sum_k \beta_{s'}^{(k)})^2\right)\left(\sum_{s<s'}\left(\alpha_{12}\beta_s^{(1)}\beta_{s'}^{(2)}\right)^2\right)}} \\
&\approx \frac{1/2\sum_{s,s'}(\sum_k \beta_s^{(k)})(\sum_k \beta_{s'}^{(k)})\left(\alpha_{12}\beta_s^{(1)}\beta_{s'}^{(2)}\right)}{\sqrt{1/2\left(\sum_s(\sum_k \beta_s^{(k)})^2\right)^2\left(1/2\sum_{s,s'}\left(\alpha_{12}\beta_s^{(1)}\beta_{s'}^{(2)}\right)^2\right)}} \\
&= \text{sign}(\alpha_{12})\frac{\left(\beta_1^T\beta_2 + \|\beta_1\|^2\right)\left(\beta_1^T\beta_2 + \|\beta_2\|^2\right)}{\left(\|\beta_1\|^2 + 2\beta_1^T\beta_2 + \|\beta_2\|^2\right)\|\beta_1\|\|\beta_2\|} \implies \\
|\gamma| &= \frac{\left(\rho h_1 h_2 + h_1^2\right)\left(\rho h_1 h_2 + h_2^2\right)}{\left(h_1^2 + 2\rho h_1 h_2 + h_2^2\right)h_1 h_2}
\end{aligned}$$

We define $\rho = \widehat{\text{Cov}}(\beta^{(1)}, \beta^{(2)}) = \beta_1^T\beta_2$ (informally identifying $\beta_i$ and $\beta^{(i)}$ where convenient). Because we assumed only $K = 2$ pathways, the space of inter-pathway correlation collapses to the scalar $\rho$. We assume $\rho \geq 0$ for simplicity and because this is usually biologically plausible, but could do similar calculations for $\rho < 0$.

Letting $x := h_2/h_1$,

$$
\begin{aligned}
|\gamma| &= \frac{(\rho x + 1)(\rho + x)}{1 + 2\rho x + x^2} \\
&= \frac{\rho + (1+\rho^2)x + \rho x^2}{1 + 2\rho x + x^2} \\
&= \rho \frac{1 + 2\rho x + x^2}{1 + 2\rho x + x^2} + \frac{(1+\rho^2)x - 2\rho^2 x}{1 + 2\rho x + x^2} \\
&= \rho + (1-\rho^2)\frac{x}{1 + 2\rho x + x^2} \implies \\
|\gamma| &= \rho + (1-\rho)^2 \frac{\sqrt{h_1^2 h_2^2}}{h_1^2 + 2\rho h_1 h_2 + h_2^2}
\end{aligned} \tag{13}
$$

This exactly agrees with the above equation (10) when $\rho = 0$, as it should. On the other hand, as $\rho \to 1$, $|\gamma| \to \rho$, and in particular $\rho = 1$ gives perfect coordination. On the other end, for small $\rho$, the sign of $\gamma$ and $\rho$ can mismatch; however, this holds only when the pathways explain very different amounts of heritability and also $\rho$ is small (Figure 1). More generally, $\rho > 0$ serves to increase $\gamma$ estimates, hence showing another source of coordination.

In general, as a function of $x := h_2/h_1 \geq 0$, $\gamma$ varies over the interval $[\rho, \rho + \frac{(1-\rho)^2}{4}] = [\rho, \frac{(1+\rho)^2}{4}]$, because $4x \leq 1+2x+x^2$ for all $x \in \mathbb{R}$. We visualize these components as the upper/lower bounds in Figure 1, and illustrate how $\gamma$ varies, especially for $\rho < 0$, as a function of the ratio of heritabilities $h_1^2/h_2^2$ (i.e. holding constant total heritability).

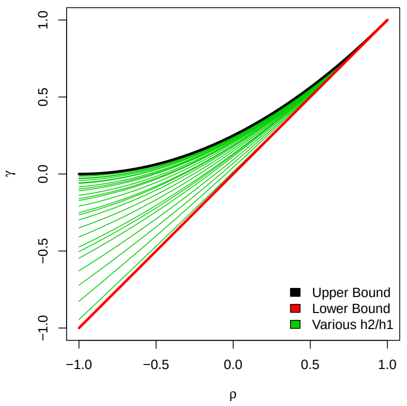

Figure 1: The range of $\gamma$ implied by different choices of $\rho$. Upper bound corresponds to $h_1^2 = h_2^2$, and lower bound corresponds to $h_1^2 \approx 0$ or $h_2^2 \approx 0$. Green lines indicate intermediate ratios $h_1^2/h_2^2 \in (0, \infty)$.

### 4 Biological models of coordinated interaction

#### 4.1 Master transcription factor

eQTLs are SNPs that effect the expression of a gene. Here, we build a phenotypic model where SNPs effect the trait only through their impact on gene expression. We partition the eQTLs into $S_0$, for a master TF, and $S_g$, for all the genes $g$ regulated by the master TF. We assume each gene is determined by the linear, *cis* effects from these eQTLs:

$$\tilde{Z}_{ig} = \mu_g + G_{i(g)}\eta_{(g)} + \delta_{ig}$$
$$Z_{i0} = \mu_0 + G_{i(0)}\eta_{(0)} + \delta_{i0}$$

Each $\eta_{(g)}$ represents the effects of *cis* SNPs on some gene, and here $\delta_{ig}$ indicates nongenetic determinants of gene expression.

Except for the master TF, we model each gene as the product of this *cis* based genetic effect and the activity level of the master TF:

$$Z_{ig} = \tilde{Z}_{ig} Z_{i0}$$

Finally, we assume the phenotype is determined by linear gene effects plus some noise term $\epsilon$:

$$y_i = \sum_{g>0} Z_{ig}\psi_g + \epsilon_i$$

Intuitively, the master TF scales the *cis*-genetic architecture of the genes it regulates, either amplifying or dampening the *cis*-signal depending on the expression of the master TF ($Z_{i0}$). In this way, the organism can control the phenotypic penetrance of entire genetic programs without altering their *cis*-genetic architecture. This resembles a *trans*-genetic regulation mechanism that was recently proposed as a biologically plausible mechanism for the omingenic model [18].

This master TF model can be recast as a pairwise SNP interaction model by expanding $Z_{ig}$:

$$\begin{aligned}
y_i &= \sum_g \left( \left(\mu_g + G_{i(g)}\eta_{(g)} + \delta_{ig}\right) \cdot \left(\mu_0 + G_{i(0)}\eta_{(0)} + \delta_{i0}\right)\right) \psi_g + \epsilon_i \\
&= \mu_0 \sum_g \mu_g\psi_g + \mu_0 \sum_g \psi_g \sum_{s\in S_g} G_{is}\eta_s + \sum_g \mu_g\psi_g \sum_{s\in S_0} G_{is}\eta_s + \sum_{g,s\in S_g;s'\in S_0} G_{is}G_{is'}\eta_s\eta_{s'}\psi_g + \epsilon'_i \\
\Longrightarrow y &= \mu' + G\beta + (G * G)\omega + \epsilon'
\end{aligned}$$

This shows our stylized master TF model falls within the framework of the interacting pathway model in (9), where:

$$\beta_s := \eta_s \begin{cases} \mu_0\psi_g & \text{if } s \in S_g \\ \sum_g \mu_g\psi_g =: w_0 & \text{if } s \in S_0 \end{cases}$$
$$\omega_{ss'} := I\{s \in S_0, s' \in S_g\}\eta_s\eta_{s'}\psi_g = \frac{I\{s \in S_0\}}{\mu_0 w_0}\beta_s\beta_{s'}$$

These definitions assume, WLOG, that SNPs are indexed such that the smallest indices all correspond to $S_0$. We also define the new noise term $\epsilon'$ by combining the phenotypic noise $\epsilon$ and the phenotypic contribution of gene expression noise in $\delta$.

Note that $\Omega$ is essentially proportional to $\beta\beta^T$, except that $\Omega_{ss'}$ is only nonzero when $s$ corresponds to the master TF and $s'$ corresponds to a regulated gene. Intuitively, then, we expect $\gamma$ to be $\pm 1$ divided by the fraction of the genome covered by *cis* SNPs for the master TF. In terms of block matrices, we are asking about the block-wise correlations between the lower triangles of $\beta\beta^T$ and $\Omega$:

$$\underbrace{\begin{pmatrix} \beta_{(1)}\beta_{(0)}^T & & & \\ \vdots & \ddots & & \\ \beta_{(S-1)}\beta_{(0)}^T & \beta_{(S-1)}\beta_{(1)}^T & \ddots & \\ \beta_{(S)}\beta_{(0)}^T & \beta_{(S)}\beta_{(1)}^T & \dots & \beta_{(S)}\beta_{(S-1)}^T \end{pmatrix}}_{\beta\beta^T} \text{ and } \underbrace{\begin{pmatrix} \beta_{(1)}\beta_{(0)}^T & & & \\ \beta_{(2)}\beta_{(0)}^T & 0 & & \\ \vdots & \vdots & \ddots & \\ \beta_{(S)}\beta_{(0)}^T & 0 & \dots & 0 \end{pmatrix}}_{\mu_0 w_0 \Omega}$$

Let $h^2_{\neg 0} := \sum_{g>0} h^2_g$ and $h^2_\beta := h^2_0 + h^2_{\neg 0}$. Visually, the correlation decomposes into column-wise terms capturing the master TF ($\sqrt{h^2_0/h^2_\beta}$) and row-wise terms capturing its downstream genes ($\sqrt{h^2_{\neg 0}/h^2_\beta}$). Then $\gamma$ can be derived directly from (10):

$$\gamma = \frac{\sqrt{h^2_0 h^2_{\neg 0}}}{h^2_0 + h^2_{\neg 0}} \tag{14}$$

Because $(a+b)^2 > 4ab$ for all $a, b \in \mathbb{R}$, $\gamma \in [-1/2, 1/2]$, consistent with our interpretation of $\gamma$ as a correlation. The factor of 2 accounts for the fact that there is no interaction between at least half of all pairs of SNPs, i.e. SNP pairs where both contribute to the master TF or to a regulated genes, and is an upper bound that is obtained only when marginal signal is evenly divided between the master TF and the combination of its regulated genes' marginal signals. These values for $h^2_0$ and $h^2_g$ also maximize the proportion of heritability due to interaction, $h^2_\omega/(h^2_\omega + h^2_\beta)$.

(14) does not exactly agree with the approximation in (4) because the latter makes an independence assumption that does not hold for this model with a single TF, essentially because it is very sparse. Nonetheless, under the further assumption that each gene explains roughly equal heritability ($h^2_0 = h^2_g = h^2_*$), then equation (14) becomes:

$$\gamma \approx \frac{\sqrt{h^2_*\left((G-1)h^2_*\right)}}{Gh^2_*} \approx \frac{1}{\sqrt{G}} = \frac{1/G}{\sqrt{(1/G)^2 + (1/G \cdot (G-1)/G)}} = \frac{\bar{\tau}}{\sqrt{\bar{\tau}^2 + \mathbb{V}(\tau)}} = (4)$$

This uses $\tau_{ss'} := \omega_{ss'}/(\beta_s\beta_{s'})$. It is not a coincidence that these equations agree under the assumption that each gene is roughly equally heritable–this assumption makes $\tau$ and $\beta \otimes \beta$ roughly uncorrelated, the condition for (4).

Finally, the AB coordination can be derived directly from (12): assume WLOG that $S_0 \in A$, because $S_0$ is assumed to be small and $S_0$, $A$ and $B$ are assumed contiguous. Also assume that $h^2_{\neg 0}$ is divided evenly across $A$ and $B$, as expected if the split is chosen randomly. Before we use (11), we must first symmetrize $\Omega$ (i.e. update $\Omega$ with $1/2(\Omega + \Omega^T)$, for the purpose of this calculation):

$$\begin{aligned}
\gamma_{AB}^2 &= \frac{1}{2}\frac{(h_0^2)(h_{\neg 0}^2/2) + (h_{\neg 0}^2/2)(0)}{(h_0^2 + h_{\neg 0}^2/2)(h_{\neg 0}^2/2)} \\
&= \gamma^2 \frac{(h_0^2 + h_{\neg 0}^2)^2}{(2h_0^2 + h_{\neg 0}^2)h_{\neg 0}^2} \\
&= \gamma^2 \left(1 + \frac{h_0^4}{2h_0^2 h_{\neg 0}^2 + h_{\neg 0}^4}\right) \\
&\approx \gamma^2
\end{aligned}$$

where the approximation holds for small $h_0^2$, i.e. when the master TF marginally explains a small fraction of overall heritability. For $h_0^2 > 0$, however, $\gamma_{AB}^2$ is slightly inflated due to the asymmetry between $A$ and $B$ that is induced by the single, strong master TF, which violates the infinitessimal model.

Note that, in this example, the master TF is the key component of the underlying biology. Nonetheless, it can explain small additive heritability relative to the rest of the genome.

#### 4.2 Polygenic buffer

The polygenic buffer model is similar to the above master TF model, in that two linear, fundamentally different systems interact. Above, each gene interacts with a single master TF; here, a polygenic buffer interacts with the individual effect of each SNP.

Partition the SNPs into $S_0$, for the polygenic buffer, and $S_1$, for the directly trait-relevant SNPs. We assume each pathway is linear and polygenic, as for the master TF, but now both pathways are built from genome-wide SNPs:

$$\begin{aligned}
Z_{i1} &= \mu_g + G_{i(1)}\eta_{(1)} + \delta_{i1} \\
Z_{i0} &= \mu_0 + G_{i(0)}\eta_{(0)} + \delta_{i0} \\
y_i &= Z_{i0} + Z_{i0} + \psi Z_{i0} Z_{i1} + \epsilon_i
\end{aligned}$$

As for the master TF model, the polygenic buffer can be recast within the polygenic model with saturated pairwise SNP interactions (9). However, the pathways now potentially contain the same SNPs, as buffer SNPs may also have trait-specific effects. Because of this genetic correlation, we use the expression for $\gamma$ derived in (13):

$$|\gamma| = \rho + (1-\rho)^2 \frac{\sqrt{h_0^2 h_1^2}}{h_0^2 + 2\rho h_0 h_1 + h_1^2}$$

where $\rho := \mathrm{Cor}(G\beta_0, G\beta_1)$ is the genetic correlation between the polygenic buffer and trait-specific genetic effects. In the case that the causal SNPs for each pathway are disjoint, or the SNP buffer effects and trait-specific effects are uncorrelated, then $\rho = 0$. In this case, the resulting $\gamma$ coincides with the master TF $\gamma$ derived in (14).

#### 4.3 Gene-environment dependence and interaction

Gene-environment interaction (GxE) can be an important component of genetic architecture across diverse organisms and biological programs. Although GxE seems fundamentally distinct from epistasis (also called GxG), the two concepts come together when the environment E is heritable. In this case, E is essentially another latent pathway in our CI model–mathematically, there is no need for the pathway to live inside the organism.

In general, we expect that practically useful choices for the environment in GxE will have some genetic basis, as is the case for most biologically meaningfully measurements. For example, smoking is an "environment" that significantly modifies the penetrance of BMI SNPs [19], and stress is another "environment" that significantly modifies the penetrance of major depression SNPs [20]. But smoking and stress are clearly heritable, and are themselves often treated as phenotypes of direct interest.

Mathematically, we assume a GxE model where the polygenic score $G\beta_g$ interacts with the continuous environment $e'$:

$$y = G\beta_g + e' + \zeta(G\beta_g) \circ e' + \epsilon'$$

We also allow that the environment may have some genetic basis: $e' = G\beta_e + e$. This gives:

$$\begin{aligned} y &= G(\beta_g + \beta_e) + e + \zeta(G\beta_g) \circ (G\beta_e) + \zeta(G\beta_g) \circ e + \epsilon' \\ &=: G(\beta_g + \beta_e) + (G * G)(\zeta\beta_g \otimes \beta_e) + \epsilon \end{aligned}$$

This collapses $e$ and $(G\beta_g) \circ e$ into $\epsilon$. As $e$ is independent of $G$, $\epsilon$ will be roughly Gaussian if $\epsilon'$ is roughly Gaussian.

This is now in the form of the interacting pathways model in (9). However, the pathways may now be correlated if there is genetic correlation $\rho$ between the direct effect of $G$ on the trait ($\beta_g$) and the direct effect of $G$ on the environment ($\beta_e$). We note, however, that $\rho \neq 0$ is not implied by $h_e^2 \neq 0$, i.e. G-E correlation is insufficient to cause $\rho \neq 0$.

Using the formulas that allow inter-pathway correlation (Section 3.3), the coordination is:

$$\gamma = \text{sign}(\zeta) \cdot \left( \rho + \frac{(1-\rho)^2}{4} \frac{\sqrt{h_1^2 h_2^2}}{(1/2(h_1 + h_2))^2} \right)$$

This decomposes the coordination into two terms:

1. $\rho$, which captures the coordination from correlated pleiotropic effects of $G$ on $y$ and $e'$.
2. $\frac{(1-\rho)^2}{4} \frac{\sqrt{h_g^2 h_e^2}}{(1/2(h_g+h_e))^2}$, which captures the coordination from the interaction between the direct genetic pathway and the genetic pathway through the environment $e'$. The deflation factor $\frac{(1-\rho)^2}{4}$ (cf. equation (10)) adjusts for double-counting the overlapping genetic signal between $\beta_g$ and $\beta_e$–already counted in the first term–and ensures that $|\gamma| \leq 1$.

#### 4.4 Non-sparse random effect model for $\Omega$

In this section, we assume that $\Omega$ has a random effect distribution.

$$\omega_{ss'} | \beta \overset{\text{ind}}{\sim} \mathcal{N}\left(\tau\beta_s\beta_{s'}, \sigma_\omega^2 \sigma_\beta^4\right) \tag{15}$$

This model on $\Omega$ is identical to previous work for genome-by-genome interactions when $\tau = 0$, as coordination drops out [2–10].

First, we assume that $S$ is large enough so that:

$$\gamma = \frac{\sum_{s>s'} \Omega_{ss'}\beta_s\beta_{s'}}{\sqrt{\sum_{s>s'} \Omega_{ss'}^2}\sqrt{\sum_{s>s'} \beta_s^2\beta_{s'}^2}} \approx \frac{\tau}{\sqrt{\tau^2+\sigma_\omega^2}}$$

Write $\Omega = \tau\beta\beta^T + \Omega'$, and define $h_A^2 := \frac{1}{|A|}\sum_{s\in A}\beta_s^2$ and $h_B^2 := \frac{1}{|B|}\sum_{s\in B}\beta_s^2$. The $AB$ coordination is:

$$\begin{aligned}
\gamma_{AB} &= \frac{\beta_A^T\Omega_{AB}\beta_B}{\sqrt{h_A^2h_B^2h_{AB}^2}} \\
&= \frac{\tau\sum_{s\in A,s'\in B}\beta_s^2\beta_{s'}^2 + \sum_{s\in A,s'\in B}\Omega'_{ss'}\beta_s\beta_{s'}}{\sqrt{\sum_{s\in A,s'\in B}\left(\tau^2\beta_s^2\beta_{s'}^2 + {\Omega'_{ss'}}^2\right)}\sqrt{\sum_{s\in A,s'\in B}\beta_s^2\beta_{s'}^2}} \\
&\approx \frac{\tau h_A^2h_B^2}{\sqrt{h_A^2h_B^2\left(\tau^2h_A^2h_B^2 + |A||B|\sigma_\omega^2\sigma_\beta^4\right)}} + \delta_1 \\
&= \frac{\tau(\sigma_\beta^2|A|)(\sigma_\beta^2|B|)}{\sqrt{(\sigma_\beta^2|A|)(\sigma_\beta^2|B|)\left(\tau^2(\sigma_\beta^2|A|)(\sigma_\beta^2|B|) + |A||B|\sigma_\omega^2\sigma_\beta^4\right)}} + \delta_1 + \delta_2 \\
&= \frac{\tau}{\sqrt{\tau^2+\sigma_\omega^2}} + \delta_1 + \delta_2 \\
&\approx \gamma
\end{aligned}$$

The first approximation assumes that $S$ is large enough so that, for any $A, B$, it approximately holds that $\frac{1}{|A||B|}\sum_{s\in A,s'\in B}{\Omega'_{ss'}}^2 \approx \mathbb{V}(\Omega_{ss'}) = \sigma_\omega^2\sigma_\beta^4$. The second approximation assumes that $\delta_1$ and $\delta_2$ are negligible, which we motivate below.

The $\delta_1$ term is just mean zero noise deriving from random uncoordinated fluctuations in $\Omega$. This term vanishes as the number of interacting SNPs, $|A||B|$, increases, as it averages out over the increasingly large number of comparisons. More formally:

$$\delta_1 := \frac{\sum_{s\in A,s'\in B}\Omega'_{ss'}\beta_s\beta_{s'}}{\sqrt{h_A^2h_B^2\left(\tau^2h_A^2h_B^2 + |A||B|\sigma_\omega^2\sigma_\beta^4\right)}} \implies \delta_1|A,B,\beta \sim \mathcal{N}\left(0, \frac{\sigma_\omega^2}{\tau^2h_A^2h_B^2 + |A||B|\sigma_\omega^2\sigma_\beta^4}\right)$$

In particular, this shows that for all $A$ and $B$ we have $\mathbb{V}(\delta_1|A,B,\beta) \leq \frac{1}{|A||B|\sigma_\beta^4}$, hence $\delta_1 \to 0$ deterministically as $|A||B|$ grows.

For $\delta_2$, let $\varepsilon := \frac{h_A^2 h_B^2}{|A||B|\sigma_\beta^4} - 1$. Then:

$$\begin{aligned}
\delta_2 &:= \frac{\tau\sqrt{h_A^2 h_B^2}}{\sqrt{\tau^2 h_A^2 h_B^2 + |A||B|\sigma_\omega^2\sigma_\beta^4}} - \frac{\tau}{\sqrt{\tau^2+\sigma_\omega^2}} \\
&= \sqrt{\frac{\varepsilon+1}{\varepsilon+1+\sigma_\omega^2/\tau^2}} - \frac{\tau}{\sqrt{\tau^2+\sigma_\omega^2}} \\
&\approx \left(\left(\sqrt{\frac{1}{1+\sigma_\omega^2/\tau^2}}\right) + \varepsilon\left(-\frac{\sigma_\omega^2}{2(\tau^2+\sigma_\omega^2)}\right)\right) - \frac{\tau}{\sqrt{\tau^2+\sigma_\omega^2}} \\
&= -\frac{\sigma_\omega^2}{2(\tau^2+\sigma_\omega^2)}\varepsilon
\end{aligned}$$

The approximation is a first-order approximation of $\varepsilon$ around 0, i.e. assuming $h_A^2 h_B^2 \approx |A||B|\sigma_\beta^4$. Essentially, this approximation assumes that CI is stable over most random $A/B$ splits, as expected for polygenic models.

We do not pursue formally characterizing the distribution of $\varepsilon$ as a function of random $A$ and $B$. But we list a few facts to demonstrate that the approximation $\varepsilon \approx 0$ is reasonable for polygenic models:

- By the LLN, as $|A|, |B| \to \infty$, then $h_A^2 h_B^2 \to \sigma_\beta^4 |A|B| \implies \varepsilon \to 0$.
- By the CLT, $h_A^2 \to \mathcal{N}\left(\sigma_\beta^2|A|, \frac{S}{|A|/S(1-|A|/S)}\sigma_\beta^2\right)$ in distribution as the number of (causal) SNPs in $A$ grows. The variance calculation assumes $S$ is large enough to approximate the probability of SNPs being in $A$ or $B$ as independent across SNPs.
- Assuming $h_A^2 \sim \mathcal{N}\left(\sigma_\beta^2|A|, \frac{S}{|A|/S(1-|A|/S)}\sigma_\beta^2\right)$, then:

$$\mathbb{V}(\varepsilon) = \frac{1}{S-|A|}\left(\frac{S}{|A|} - 4\frac{|A|}{S} - 2\frac{1}{|A|S\sigma_\beta^4}\right) \to 0$$

  where the convergence is for $|A|, S \to \infty$.

### 5 Technical results

#### 5.1 $\hat{\gamma}_{AB}$ is consistent and unbiased for $\gamma_{AB}$

**Proposition 1.** *Assume the linear model in* (1)*, and that the $S$ SNPs are partitioned into "independent" subsets $A$ and $B$, i.e.:*

$$r^2(A,B) := \max_{s\in A, s'\in B} \frac{1}{N}\sum_{i=1}^N G_{is}G_{is'} = 0$$

*Assume also that $G$ has columns scaled to mean zero and variance one.*

*Define the A/B coordination as*

$$\gamma_{AB} = \frac{\beta_A^T \Omega_{AB} \beta_B}{\|\beta_A\| \|\beta_B\| \|\Omega_{AB}\|} \tag{16}$$

*then the even/odd estimator of* $\gamma$ *w.r.t. the partition* $A$ *and* $B$ *defined in* (7) *is:*

$$\hat{\gamma}_{AB} = \gamma_{AB} + \mathcal{N}\left(0, \frac{1}{N\|\Omega_{AB}\|^2}\right) \tag{17}$$

**Proof**

We are interested in the OLS estimate of $\gamma$. Letting $\Pi^{\perp}$ be the orthogonal projection onto $P := (P_A|P_B)$, the two stage least squares expression for the OLS estimate of $\hat{\gamma}$ is:

$$\hat{\gamma} := \frac{[P_A \circ P_B]^T \, \Pi^{\perp} y}{[P_A \circ P_B]^T \, \Pi^{\perp} \, [P_A \circ P_B]}$$

We assume that $P_A$ and $P_B$ are created from independent SNPs and that sample size is sufficiently large to replace empirical moments by their theoretical expectations. This implies that the PRS interaction terms $P_A \circ P_B$ orthogonal to both $P_A$ and $P_B$, which in turn implies the OLS estimate of $\lambda$ is:

$$\hat{\lambda}_{OLS} \approx \frac{[P_A \circ P_B]^T \left([G * G]\,\omega + \epsilon\right)}{[P_A \circ P_B]^T \, [P_A \circ P_B]}$$

Expanding these product in the denominator gives:

$$\begin{aligned}
[P_A \circ P_B]^T \, [P_A \circ P_B] &= \sum_i \sum_{s,j \in A, s', j' \in B} \left(G_{is} G_{ij} G_{is'} G_{ij'}\right) \beta_s \beta_j \beta_{s'} \beta_{j'} \\
&= \sum_{s,j \in A, s', j' \in B} \left(\sum_i G_{is} G_{ij} G_{is'} G_{ij'}\right) \beta_s \beta_j \beta_{s'} \beta_{j'} \\
&= \sum_{s,j \in A, s', j' \in B} \left(N \left[\sum_i G_{is} G_{is'}\right] \left[\sum_i G_{ij} G_{ij'}\right]\right) \beta_s \beta_j \beta_{s'} \beta_{j'} && (\dagger) \\
&= N \beta_A^T R_A \beta_A \beta_B^T R_2 \beta_B \\
&= N \|\beta_A\|^2 \|\beta_B\|^2 && (\ddagger)
\end{aligned}$$

† uses the assumption of no LD between $A$ and $B$, and ‡ uses the much stronger assumption that all causal SNPs are in linkage equilibrium. A similar expansion in the numerator gives:

$$\begin{aligned}
\hat{\lambda} &= \frac{\sum_{s \in A, s' \in B, j, j', i} \left(G_{is} G_{ij} G_{is'} G_{ij'}\right) \beta_s \beta_{s'} \omega_{jj'} + \sum_{s \in A, s' \in B} \left(G_{is} G_{is'} \beta_s \beta_{s'} \epsilon_i\right)}{N \|\beta_A\|^2 \|\beta_B\|^2} \\
&\stackrel{d}{=} \frac{\beta_A^T \Omega_{AB} \beta_B}{\|\beta_A\|^2 \|\beta_B\|^2} + \mathcal{N}\left(0, \frac{1}{N\|\beta_A\|^2 \|\beta_B\|^2}\right) \\
&\stackrel{d}{=} \gamma_{AB} \frac{\|\Omega_{AB}\|}{\|\beta_A\| \|\beta_B\|} + \mathcal{N}\left(0, \frac{1}{N h_A^2 h_B^2}\right) \implies \\
\hat{\gamma}_{AB} &\sim \gamma_{AB} + \mathcal{N}\left(0, \frac{1}{N\|\Omega_{AB}\|^2}\right)
\end{aligned}$$

#### 5.2 Monotone phenotype transformations

We consider the effect of nonlinear phenotype transformations on coordination. Assume that trait values are transformed by some twice-differentiable function $f : \mathbb{R} \mapsto \mathbb{R}$. We assume that $f$ is monotone, which is equivalent to the notion of rescaling $y$ because the order of the traits $f(y_i)$ is the same as the order of the traits $y_i$.

The coordination definition given in (3) assumes that the pairwise interaction model in (1) holds exactly for some true $\beta$ and $\Omega$. However, if $f$ is applied to each phenotype value, then clearly (1) no longer holds for the transformed phenotype; moreover, unless $f$ is linear, (1) does not hold for *any* values of $\beta$ and $\Omega$. Hence it is not clear how to define $\gamma$ by (3).

Instead, we define the coordination under the transformation $f$ as the coordination amongst the least-squares best fits for the linear and interaction parameters, $\tilde{\beta}$ and $\tilde{\Omega}$:

$$\gamma_f(\Omega, \beta) := \gamma(\tilde{\Omega}, \tilde{\beta}) \tag{18}$$

We assume perfect coordination on the original scale for ease, so $y := g\beta + \lambda\,(g \otimes g)\,(\beta \otimes \beta) + \epsilon$. Then the least-squares estimates at some reference genotype value $g \in \mathbb{R}^S$ are given by:

$$\begin{aligned}
\nabla_{g_s} f(y) &= f'(y)\beta_s(1 + \lambda g\beta) \\
\nabla^2_{g_s, g_{s'}} f(y) &= f''(y)\beta_s\beta_{s'}(1 + \lambda g\beta)^2 + f'(y)(\lambda\beta_s\beta_{s'}) \\
&= \beta_s\beta_{s'}\left(f''(y)(1 + \lambda g\beta)^2 + \lambda f'(y)\right) \\
\implies \gamma_f(g) &:= \mathrm{Cor}\left((\nabla f)_g \otimes (\nabla f)_g, (\nabla^2 f)_g\right) \\
&= \mathrm{Cor}\left((f'(y)\beta(1 + \lambda g\beta)) \otimes (f'(y)\beta(1 + \lambda g\beta)), \beta \otimes \beta\left(f''(y)(1 + \lambda g\beta)^2 + \lambda f'(y)\right)\right) \\
&= \mathrm{sign}\left(f''(y)(1 + \lambda g\beta)^2 + \lambda f'(y)\right)
\end{aligned}$$

When $|f'\lambda|/|f''|$ is large, the second term dominates and

$$\gamma_f(g) = \mathrm{sign}(\lambda f'(y)) = \gamma\,\mathrm{sign}(f') \tag{19}$$

The last equality using the fact that $\gamma = \mathrm{sign}(\lambda)$ and emphasizes that $\mathrm{sign}(f')$ is constant for all $y$ because $f$ is monotone. (19) is the "right" answer, in the sense that $\gamma$ is unchanged except, perhaps, a sign flip if $f$ is decreasing.

On the other hand, if $|f'|/|f''|$ is small–meaning the function varies wildly compared to linear–or if $|\lambda|$ is small–meaning the overall interaction signal is weak–then $\gamma_f(g)$ may have the incorrect sign.

On average over mean-zero genotypes $g$, the former force wins out, showing coordination is on balance preserved under relatively smooth, strictly monotone functions. Assume that $\lambda > 0$ (a symmetric argument applies for $\lambda < 0$), so:

$$\begin{aligned}
\mathbb{E}\left(f''(y)(1 + \lambda g\beta)^2 + \lambda f'(y)\right) &\geq \mathbb{E}\left(-\|f''\|_\infty(1 + \lambda g\beta)^2\right) + \lambda\mathbb{E}\left(f'(y)\right) \\
&= -\|f''\|_\infty(1 + \lambda^2\sigma_g^2) + \lambda\bar{f}' \geq 0 \iff \\
\mathrm{sign}(f')\frac{\lambda}{(1 + \lambda^2\sigma_g^2)} &\geq \frac{\|f''\|_\infty}{|\bar{f}'|}
\end{aligned}$$

This has the same sign as $\lambda$ so long as $|f''/\bar{f}'|$ never grows too large (over the distribution of $g$), i.e. the function never curves dramatically relative to its overall linear approximation. Large is defined in comparison to $\lambda$, which is basically (proportional to) the left hand side for realistic parameters.

### 6 Simulation details

#### 6.1 Genotypes

Genotypes were generated under three scenarios: 1) absence of assortative mating and structure, 2) presence of assortative mating and absence of structure, and 3) absence of assortative mating and presence of structure. To simulate the absence of assortative mating and structure, genotypes were generated for 2,000 individuals at 500 SNPs. Genotypes for each individual $g_{ij}$ were drawn from a binomial distribution $g_{ij} \overset{\text{ind}}{\sim} \text{Binomial}(2,p_j)$, with $p_j$ drawn i.i.d. from a Beta(5,5) distribution.

To simulate the scenario of assortative mating in the absence of population structure, genotypes were generated for a parent generation of 4,000 individuals at 500 SNPs in the same manner described above for $g_{ij}$. Phenotypes for each individual in the parent generation were then calculated (details below). Individuals were sorted by phenotype and paired in sequential groups of two to simulate one generation of extreme assortative mating. Child genotypes were then generated by randomly drawing the value of one maternal allele and one paternal allele.

To simulate the scenario of population structure, genotypes were simulated for two populations separately, each with 1000 individuals and 500 SNPs, where $F_{ST} = 0.1$. Population specific allele frequencies $p_{kj}$ were drawn i.i.d. over SNPs and populations from $\text{Beta}\left(\frac{1}{2}(1 - F_{ST}), \frac{1}{2}\left(\frac{(1-F_{ST})}{F_{ST}}\right)\right)$. Each individual's genotypes were then drawn independently from Binomial$(2,p_{kj})$, where $k$ indicates each person's respective population and $j$ indicates SNPs.

##### 6.1.1 Phenotypes

For each set of simulated genotypes, phenotypes were simulated under three causal scenarios: additive, uncoordinated interaction, and coordinated interaction. To simulate phenotypes under the additive model, SNP effects $\beta$ were drawn i.i.d. from a standard normal distribution. Phenotypes for each individual were then simulated by summing genotypes $g_{ij}$ weighted by their respective effects, $\beta_j$, across all SNPs and adding a random error drawn i.i.d from a standard normal distribution.

To simulate phenotypes under uncoordinated interactions, effects for 1% of all SNP pairs were chosen to have an interaction effect $\beta_{jj'}$ which is drawn i.i.d. standard normal. Genotypes from the resulting 2,500 randomly sampled pairs of simulated SNPs were multiplied together element-wise and subsequently multiplied by the corresponding effect for each pair. These values were then summed across all 2,500 causal SNP pairs, and i.i.d. standard normal noise is added to each individual.

To simulate phenotypes under the coordinated interaction model, 200 SNPs were randomly chosen to be modified (modified group) by the effects of an independent set of 100 SNPs (buffer group). Effect sizes for all chosen SNPs in each group were equal in distribution to those in the additive model. Genotypes in the buffer group were multiplied by their respective effect sizes and then summed to create the buffer values $b$. Effect sizes of the modified group were then multiplied by 70 percent of the effect of the buffer SNPs. Genotypes in the modified group were then multiplied by the respective modified effect sizes. For each individual, phenotypes were the sum across all SNPs in the modified group with the addition of random error drawn i.i.d. from a standard normal distribution.
